# Single-Molecule Reaction-Diffusion

**DOI:** 10.1101/2023.09.05.556378

**Authors:** Lance W.Q. Xu, Sina Jazani, Zeliha Kilic, Steve Pressé

**Author notes:** Corresponding author. Email address Website: https://cbp.asu.edu/content/steve-presse-lab.

## Abstract

We propose to capture reaction-diffusion on a molecule-by-molecule basis from the fastest acquirable timescale, namely individual photon arrivals. We illustrate our method on intrinsically disordered human proteins, the linker histone H1.0 as well as its chaperone prothymosin *α*, as these diffuse through an illuminated confocal spot and interact forming larger ternary complexes on millisecond timescales. Most importantly, single-molecule reaction-diffusion, smRD, reveals single molecule properties without trapping or otherwise confining molecules to surfaces. We achieve smRD within a Bayesian paradigm and term our method Bayes-smRD. Bayes-smRD is further free of the average, bulk, results inherent to the analysis of long photon arrival traces by fluorescence correlation spectroscopy. In learning from thousands of photon arrivals continuous spatial positions and discrete conformational and photophysical state changes, Bayes-smRD estimates kinetic parameters on a molecule-by-molecule basis with two to three orders of magnitude less data than tools such as fluorescence correlation spectroscopy thereby also dramatically reducing sample photodamage.

## 1 Introduction

Individual biochemical reactions are the basis of information communication and signaling in cells [1–3]. As such, developing methods to track and study individual biochemical reactions is the key towards unraveling, on a molecule-by-molecule basis, the mechanisms of life. In order to capture individual molecular interactions at the fastest possible acquirable timescale, one may analyze photon arrivals derived from single molecules as these traverse a confocal spot [4, 5].

Typical methods of analysis of confocal data, such as fluorescence correlation spectroscopy (FCS), indeed analyze photon arrival data [6, 7] and draw conclusions on events at or below microsecond timescales [8–10]. However, in order to derive diffusion coefficients [11, 12] as well as other dynamical quantities [11–13], correlative methods rely on averaging over multiple (often thousands and more) single-molecule traversals through the confocal spot (termed bursts, see Fig. 1) [6–13]. In doing so, correlative methods provide bulk, averaged, properties. Yet information at the single-molecule level is encoded in individual bursts. This includes information on the heterogeneity of pairwise interactions of interest here, for example, the interactions of intrinsically disordered protein (IDP) pairs [14].

**Figure 1:**
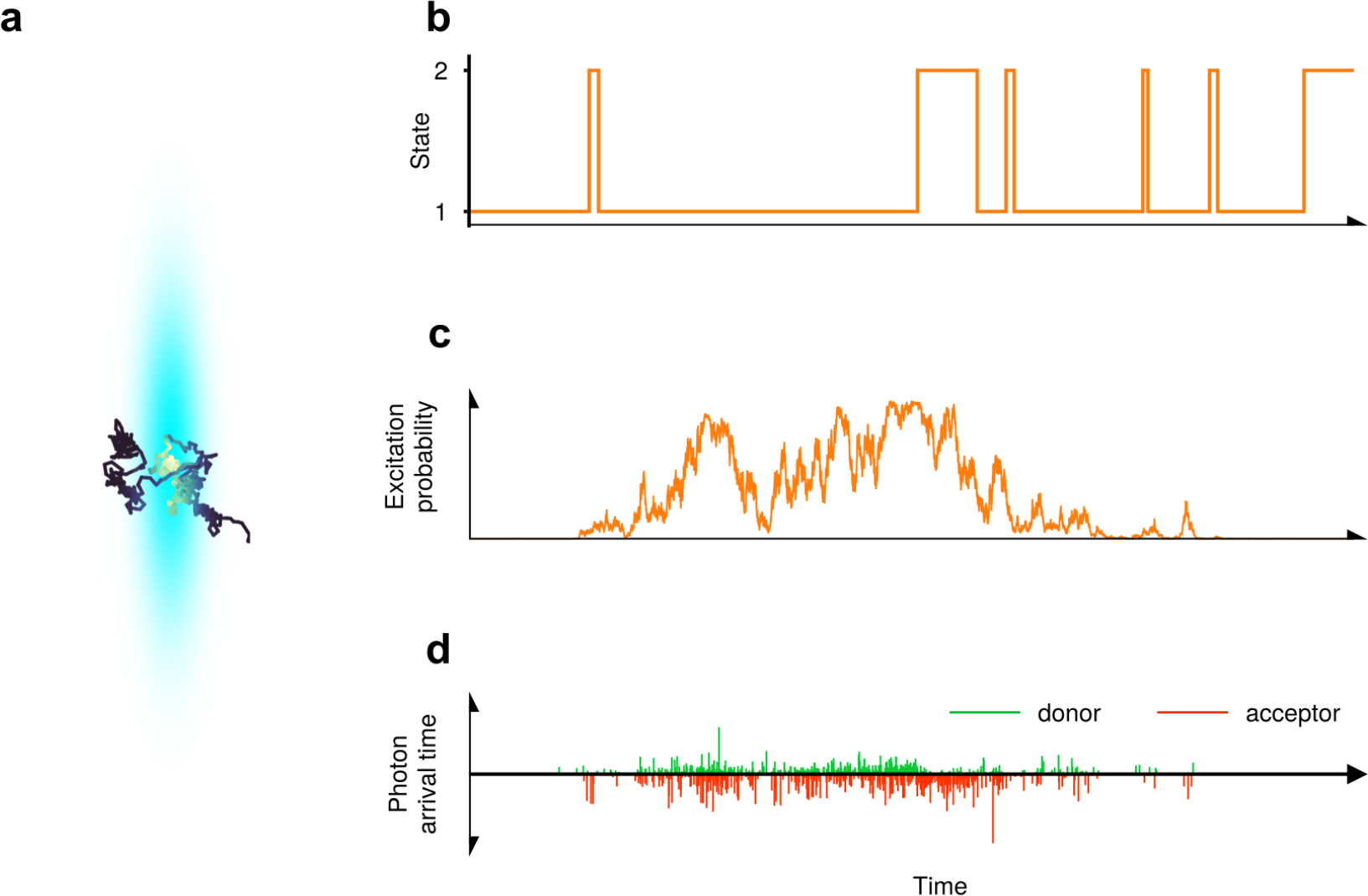
An illustration of a molecule labeled with a FRET pair undergoing state transitions while traversing a confocal spot. (a) The molecule’s 3D spatial trajectory. Darker colors indicate the molecule being further away from the spot center. (b) The molecule’s state trajectory. State 1(2) corresponds to the low(high)-FRET efficiency state. State 1 also has a slower diffusion coefficient. (c) The molecule’s probability of being excited by a laser pulse versus time. This trajectory encodes information on the excitation rate and spatial distance with respect to the confocal spot’s center. (d) A photon detection trace coming from these trajectories. A region with dense photon detections is often referred to as a burst.

Toward resolving full reaction-diffusion dynamics at the single-molecule level, attempts have been made towards obtaining reaction rates with Förster resonance energy transfer (FRET) pairs [15] as molecules diffuse through confocal volumes, although these methods lack the ability to simultaneously resolve diffusion dynamics [5, 16–18]. Other tools exist for tracking molecules through confocal spots in the absence of reaction kinetics [19–23]. Similarly, numerous methods exist for extracting FRET state transitions albeit for complexes tied to surfaces [24–28]. Yet tying substrates to surfaces can lead to changes in conformation and reactivity [29, 30] and this motivates our ambition to investigate single-molecule reaction-diffusion, smRD, for freely diffusing substrates.

In other words, we seek a method capable of extracting both reaction rates and diffusion dynamics at the single-molecule level from the fastest acquirable timescale, namely photon arrivals. As such, we propose smRD dynamics using the following information: (i) photon arrival times (*i*.*e*., excited state lifetimes) for pulses resulting in photon arrivals; (ii) whether pulses yield detectable photons (*i*.*e*., which pulses are “empty”); and (iii) donor or acceptor channels from which each photon is detected.

To avoid biases in our estimates for smRD parameters, we must also incorporate factors including background photon shot noise, fluorophore photophysical transitions (such as photoblinking and non-radiative decays), the shape of instrumental response functions (IRFs), direct acceptor excitation probability, and the bleed-through between donor and acceptor detection channels.

Given that we aim to capture both reaction kinetics and diffusion dynamics from single bursts, accounting for the factors highlighted above, our smRD framework, Bayes-smRD, must efficiently extract information from every photon. In order to achieve this goal, we operate within the Bayesian paradigm [19–21, 31–33] from which we build a hierarchical mathematical model that establishes probabilistic connections between the reaction-diffusion dynamics, and experimental observations. As all temporal correlations in the data are leveraged within this mathematical model without data pre-processing, our framework is highly efficient.

Concretely, in the Bayesian paradigm, our objective is to calculate complete probability distributions for the trajectories of the molecule states (including conformational, photophysical, and spatial) as well as their coinciding dynamical parameters (transition kinetics and diffusion coefficients) self-consistently and simultaneously. In this study, we consider discrete conformational and photophysical states, which directly influence FRET efficiency. We also consider continuous spatial states unrelated to FRET efficiency. For the sake of clarity, we use the term “state” to refer specifically to the joint conformational and photophysical state, while we use “spatial position” or “spatial trajectories” instead of speaking of a continuous spatial state.

An important concern when analyzing single photon arrivals is the possibility that the duration of a traversal, a “burst”, determined by the confocal volume size and diffusion coefficient, might be shorter than the state lifetimes. In such cases, the analyzed burst may not capture any transitions. To address this issue, our framework also allows for the analysis of multiple bursts with independent state and spatial trajectories, while assuming they share the same reaction-diffusion parameter values. Despite considering multiple bursts, the amount of data analyzed is still two to three orders of magnitude less than data typically analyzed in Ref. [34, 35]. This is a critical advantage of the method we propose in avoiding photodamage with the potential to probe reactions of photosensitive biomolecules [36, 37].

In the subsequent sections, we demonstrate Bayes-smRD on both synthetic and experimental data involving the necessarily heterogeneous pairwise interaction of intrinsically disordered human proteins prothymosin *α* (ProT*α*) to linker histone H1.0 (H1) at a single-molecule level.

## 2 Results

Within the Bayesian paradigm, we compute full joint probability distributions (termed posteriors) over a molecule’s state trajectories, spatial trajectories, state transition rates, diffusion coefficients, as well as other quantities of interest (including FRET efficiencies and excitation rate). The breadth of these posterior probability distributions, especially critical in single-molecule settings, reflects uncertainty propagated from the finiteness of the data available (as molecules come in and out of the confocal volume), but also other experimental parameters (background noise, breadth of IRF, and detector bleed-through). All posterior distributions, shown as histograms, in this paper are normalized as probability densities (with unit area). In addition to distributions, specifically for state trajectories and spatial trajectories, we also provide the *maximum a posteriori* (MAP) trajectories as point estimates. In this work, we make the assumption (though not required) of a 3D Gaussian confocal volume [38, 39]. The framework, as discussed in the Methods section, can accommodate any pre-calibrated (known) confocal volume shape. The confocal volume’s spatial symmetries result in multiple spatial trajectories sharing the same probability density, leading to degeneracy in a molecule’s absolute spatial positions. While this degeneracy does not pose a problem during the inference step, it introduces ambiguity visualizing results. Therefore, only when visualizing results, we represent the molecule’s “excitation probability trajectory” instead of its absolute spatial trajectory. The excitation probability trajectory, defined in Eq. (S.5), combines statistically equivalent spatial trajectories and provides a combined measure of the molecule’s excitation rate and its distance from the confocal spot center (see Fig. 1c).

Along these same lines, in what follows we report escape rates (inverse lifetime) of each state reflecting how fast the system leaves its current state as an alternative to reporting concentration dependent association rates otherwise less meaningful at the single-molecule level.

In the next two subsections, we first apply Bayes-smRD to experimental data collected on the interaction of IDP fragments ProT*α* and H1, then validate our framework’s robustness using synthetic data where ground truth is accessible for sake of comparison.

### 2.1 Experimental data and analysis description

In the experimental analysis, we monitor the interactions of human proteins ProT*α* (net charge -44) and linker histone H1 (net charge +53). As both are highly and oppositely charged, ProT*α* and H1 appear in bound states while retaining disordered features [40–42], as illustrated in Fig. 2. Experimental data from single-molecule, circular dichroism, and nuclear magnetic resonance spectroscopy, as well as molecular simulations [40–42] support the hypothesis of non-specific ProT*α* and H1 binding with expected diffusion-limited and concentration dependent association and rapid concentration-independent dissociation, despite their high and opposite charges [40–42]. It has been hypothesized that this type of (dis)association may help retain regulatory cell network responsiveness [41].

**Figure 2:**
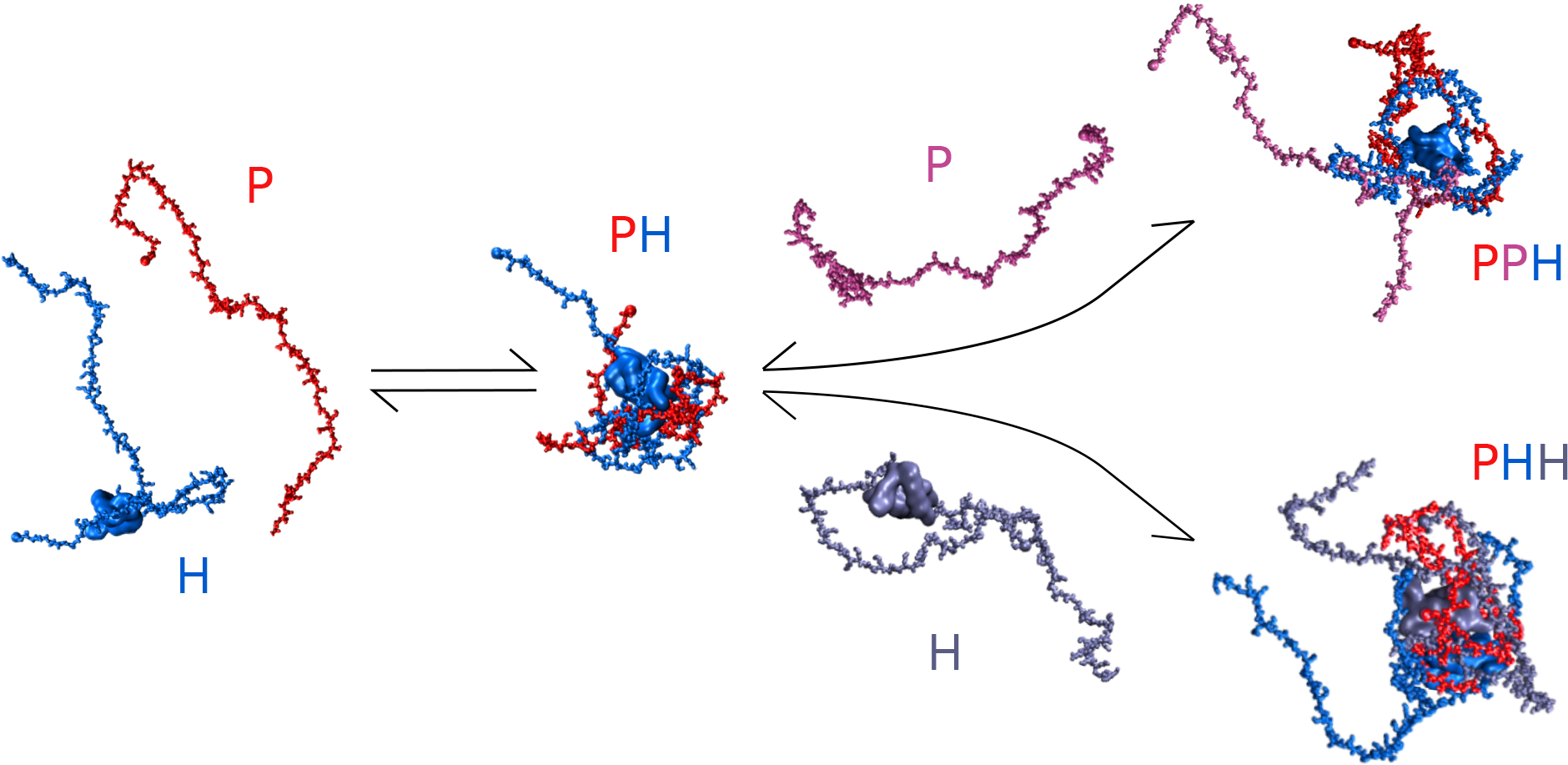
Some kinetic schemes involving ProT*α* and H1 molecules (P and H in this figure, respectively) shown with snapshots from coarse-grained molecular dynamics simulations [40, 41]. A freely diffusing ProT*α* and an H1 may form a ProT*α*-H1 (PH) dimer, compactifying their conformation. This dimer may putatively continue binding other ProT*α*’s and H1’s if available forming larger ternary complexes including ProT*α*_2_-H1 and ProT*α*-H1_2_ (PPH and PHH in this figure, respectively). Possible further reactions, *e*.*g*., the formation of tetramers, are excluded herein. At each step, this figure illustrates only one of many candidate structures for this highly dynamic and intrinsically disordered system.

Here, a fluorescent FRET pair (Alexa Fluor 488 as donor and 594 as acceptor) is attached to ProT*α* at positions 56 and 110, respectively. A free doubly labeled ProT*α*, with two dyes typically far apart, is expected to coincide with a low FRET efficiency state, while bound states of the IDP pair bring the two fluorophores on ProT*α* closer together, yielding higher FRET efficiencies [40–42].

The concentration of labeled ProT*α* in the data sets analyzed is either 50 pM or 75 pM. Given the confocal volume’s dimensions (see Table S.2), the expected number of labeled ProT*α* within the volume is less than 0.05 percent at any instant. As such, a good approximation is to assume that either zero labeled ProT*α* or, at most, one is present within the illuminated region. As a result, we anticipate data with bursts of photon arrivals indicating the presence of one ProT*α* diffusing within the volume.

From Ref. [40, 41] we know that some complexes, *e*.*g*. the ProT*α*-H1 (PH) dimer, have lifetimes longer than the average duration of a burst (about 5 ms). Therefore, for all concentrations discussed in this paper, as mentioned in the Introduction, we analyze several bursts together assuming they have independent state and spatial trajectories but otherwise share the same reaction-diffusion dynamics (*e*.*g*., same diffusion coefficients for each conformational state even if each state is not visited in each trajectory). To be more specific, from a roughly eight-minute long time trace, we select 12 short time intervals in total at different times, half of which contain bursts (the burst group) with the other half containing burst-free regions (the burst-free group). Each group has its own reaction-diffusion dynamics. See Section 4.2 for more details on how we select bursts.

### 2.2 75 pM pM labeled ProT*α*

The first data set analyzed contains only 75 pM doubly labeled ProT*α*. The burst group contains six bursts that are 34 ms long in total amounting to 5,517 photon detections. From this data set, we expect to encounter only one conformational state (and thus one FRET state) though blinking of both dyes [43, 44] may introduce apparent FRET transitions. For this reason, we run a two-state model on this data set. If our model considers more states than present in the data, we find (shown in Fig. S.1) that the additional states have, as expected, much higher uncertainty (broader posterior probability distributions) associated to state trajectories, escape rates, diffusion coefficients, and FRET efficiencies, than those of the real states. Moreover, these additional states are often hardly visited. For instance, in Fig. S.1, the additional state (state 3) is visited for less than 2% of the time.

The results of our analysis are shown in Fig. 3. In particular, we show the state trajectory of one burst in Fig. 3a, the corresponding excitation probability trajectory in Fig. 3b, from the burst shown in Fig. 3c. We also obtain both states’ escape rates (equivalent to transition rates in a two-state system), diffusion coefficients, and FRET efficiencies in Figs. 3d to 3f, respectively, jointly from all six bursts analyzed, for the reason explained in Introduction. With high certainty, Fig. 3a shows that the system visits both of the states, suggesting the existence of one more state than expected. Fig. 3f provides more evidence for the existence of this extra state as it shows two very sharp and well-separated FRET efficiencies.

**Figure 3:**
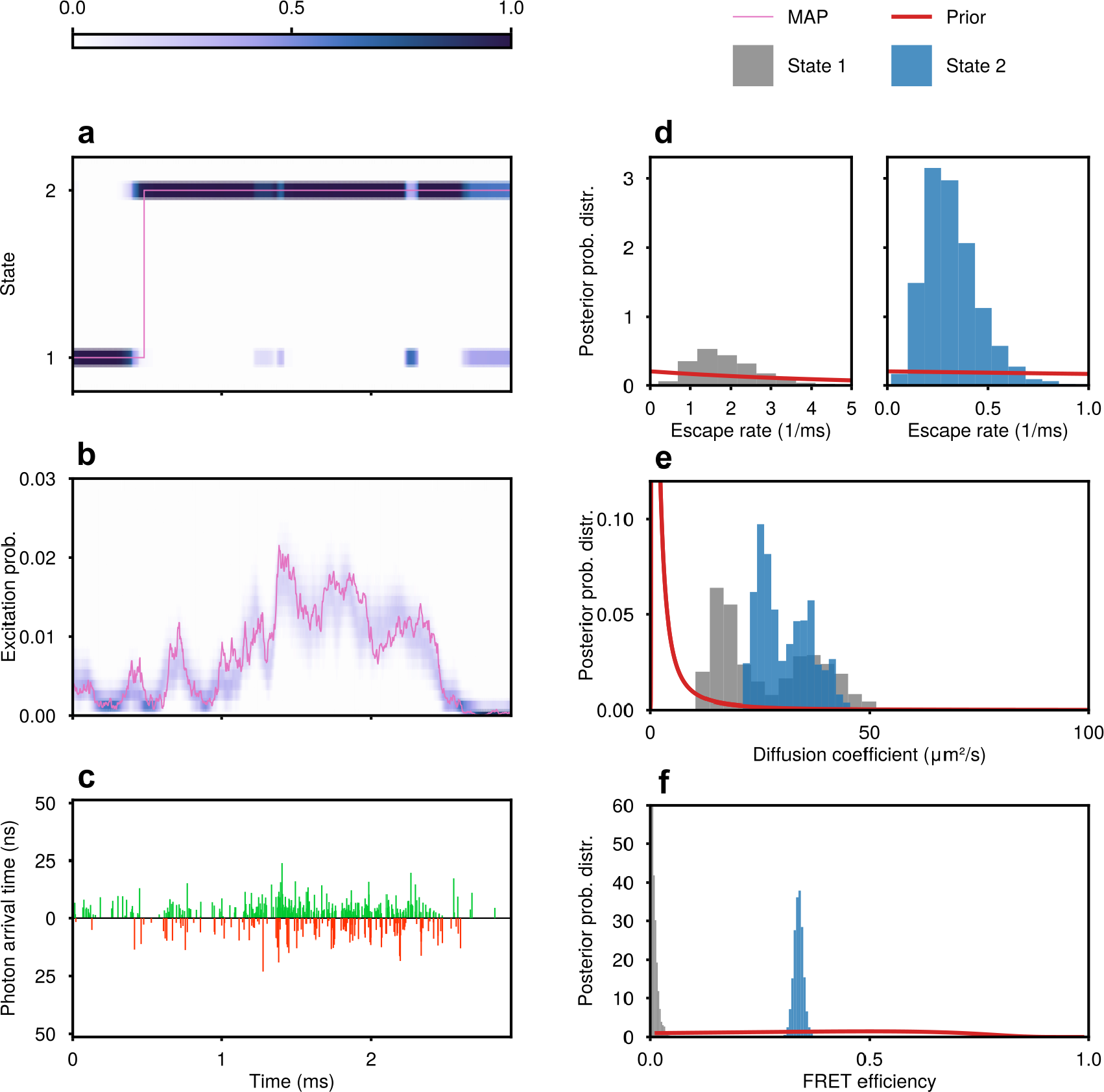
Results from the experimental data set with 75 pM labeled ProT*α* alone. (a) Learned state trajectory with the MAP trajectory highlighted by a purple line. In the main text we explain why state 1 is attributed to acceptor photophysics while state 2 corresponds to a freely diffusing labeled ProT*α*. The color map indicates confidence of the learned trajectory in terms of probability at each pulse. (b) The corresponding excitation probability trajectory which offers a joint measure of the spatial trajectory and the excitation rate. (c) The corresponding experimental data trace containing 458 detected photons within nearly 3 ms. (d)-(f) The escape rates, diffusion coefficients, and FRET efficiencies learned from six bursts (only one shown in (a)-(c)). All gray histograms are from state 1 while all blue histograms are from state 2.

One state here should coincide with the freely diffusing doubly labeled ProT*α* molecule. We attribute state 2 (blue in Figs. 3d to 3f) to this freely diffusing ProT*α* because its FRET efficiency, 0.317 to 0.359 as the 95% credible interval, is very close to that of a free ProT*α* reported in Ref. [41]. On the other hand, state 1 (gray in Figs. 3d to 3f) has zero FRET efficiency with very low uncertainty (95% credible interval is 0.000 to 0.029). This low FRET efficiency state may putatively be attributed to: (i) a conformation yielding large distance between the donor and the acceptor; (ii) free donor fluorophores in solution; and (iii) acceptor’s dark state (donor dye’s photophysics typically decreases donor channel photon counting rate and yields higher FRET efficiencies).

Here (i) is unlikely as it is not supported by any known simulation or experiment [40]. We also have reason to believe (ii), free dye, is not very likely as the 95% credible interval for the diffusion coefficient of state 1 is (12.6 to 47.4) µm^2^/s, agreeing with state 2’s diffusion coefficient 95% credible interval (22.4 to 42.0) µm^2^/s. Thus we conclude that state 1 likely originates from the acceptor’s dark state (option (iii)).

The difference in the inferred diffusion coefficient estimates presented above from those reported for free ProTα’s diffusion coefficient, 55 ±1 µm^2^/s [40], being to light an important point. Estimating diffusion coefficients using a confocal volume relies on the calibration of the volume’s shape and size. In support of our diffusion coefficient estimate, we perform FCS on the full time trace provided with more than 3*×* 10^7^ photons to learn the diffusion coefficient under the same confocal volume calibration (Table S.2). As we have argued so far, in this dataset all states share the same conformation and hence we perform a one-state FCS fit. Fig. S.2 shows our FCS curves and associated fits. The best least-square fit diffusion coefficient is 28 µm^2^/s by FCS, falling within our credible intervals. Furthermore, in Fig. S.1e where we run Bayes-smRD on the same dataset but with a three-state model, state 2’s diffusion coefficient credible interval is (21.4 to 28.8) µm^2^/s. These agreements suggest that our method learns diffusion coefficients consistently given accurate and confocal volume calibration.

Having interpreted the states and diffusion dynamics in our analysis, we now turn to the transition rates between them. As shown in Fig. 3d, the transition rate from state 2 to state 1 (*i*.*e*., the rate at which an acceptor transitions to a dark state), has 95% credible interval (0.12 to 0.64) s^−1^, while that of the reverse process is (0.67 to 4.02) s^−1^ based on all six bursts analyzed.

In order to further support the reaction-diffusion dynamics estimates reported so far, here, we also list the estimates from the same six bursts but using a three-state model, Fig. S.1. States 1 and 2 in Fig. S.1 maintain their interpretations while state 3 is intended to be redundant (and therefore has no meaningful interpretation) for the reason explained in the beginning of this section. The diffusion coefficient credible intervals of states 1 and 2 are (14.5 to 39.5) µm^2^/s and (21.4 to 28.8) µm^2^/s, respectively. As for the transition rates from 1 to 2 and from 2 to 1 in Fig. S.1, we have (0.09 to 0.77) ms^−1^ and (0.16 to 4.07) ms^−1^, respectively. Therefore, the two-state model and the three-state model yield consistent reaction-diffusion dynamics estimate.

### 2.3 75 pM labeled ProT*α* and 10 nM H1

The second data set we analyzed contains 75 pM labeled ProT*α* and 10 nM H1. As H1’s concentration is much higher than that of labeled ProT*α*, as illustrated in Fig. 2, we expect to start seeing another state corresponding to the ProT*α*-H1 complex with a higher FRET efficiency than the free ProT*α* state [40–42] as ProT*α* compactifies upon binding. At least four states are anticipated in this data set: (i) free ProT*α* with a bright acceptor, (ii) free ProT*α* with a dark acceptor, (iii) ProT*α*-H1 with a bright acceptor, and (iv) ProT*α*-H1 with a dark acceptor. Other possibilities such as a burst created by free dyes can be excluded during burst selection, see Section 4.2.

However, as states (ii) and (iv) have the same FRET efficiency (zero) and similar diffusion coefficients [40] (difference is less than the breadth of the posteriors in Fig. 3e), they are challenging to distinguish from the observables provided. Therefore, instead of treating them as separate states, we run a three-state model.

Just as before, we select a group of six bursts (5,930 photons in total) for analysis. Our results are shown in Fig. 4 following the same layout as the earlier Fig. 3. Immediately from Fig. 3f we detect a high FRET efficiency state (state 3, green) with 0.618 to 0.701, which (when comparing to the results of Fig. 3) we ascribe as originating from the ProT*α*-H1 dimer (making note that the FRET efficiency estimates depend on the pre-calibrated background photon rates, see Table S.2 for the value used in our analyses). Moreover, Fig. 3a shows that the system actually spends most amount time in state 3, consistent with Ref. [41]. As for state 3’s diffusion coefficient, Fig. 4e gives a 95% credible interval of (32.0 to 41.9) µm^2^/s. Although no literature is found to have measured this value for sake of direct comparison, Ref. [40] reports 55 ±1 µm^2^/s as free ProT*α*’s diffusion coefficient and this number decreases to 47 ±3 µm^2^/s for a mixture of ProT*α* and H1 (at a 1:1 molar ratio). Since the analyzed data set contains 75 pM labeled ProT*α* and 10 nM H1 (3:400 molar ratio), our diffusion coefficient estimate should be lower than 47± 3 µm^2^/s. This is because a higher molar ratio of H1 leads to a higher fraction of ProT*α* in its bound states (larger complexes) [41], and therefore decreases the overall diffusion coefficient. Indeed, this speculation is consistent with our estimate.

**Figure 4:**
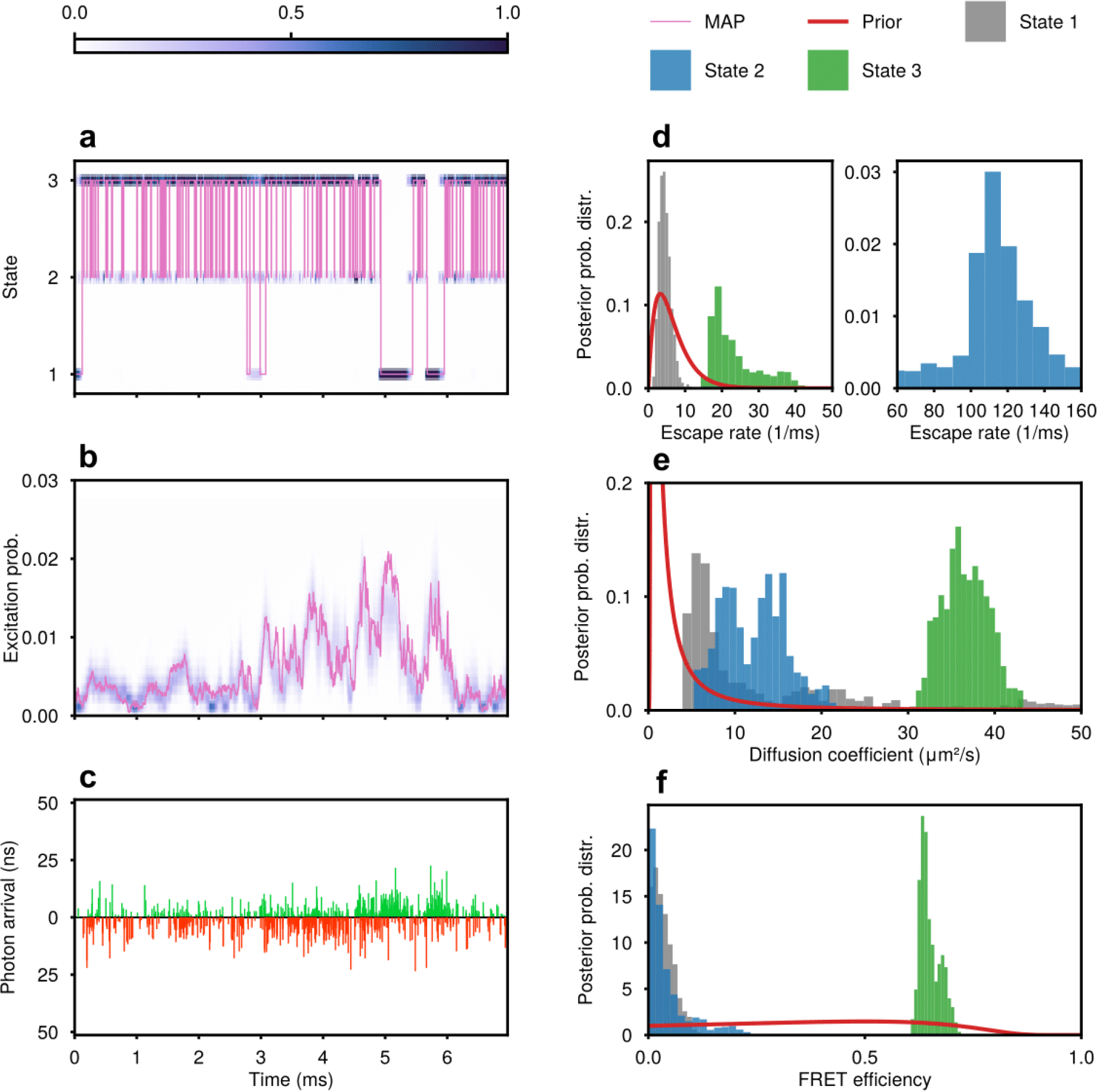
Results from the experimental data set with 75 pM labeled ProT*α* and 10 nM H1. Layout is the same as in Fig. 3. We discuss in the main body why state 3 is most likely attributed to the ProT*α*-H1 complex.

State 1 (gray), on the other hand, has FRET efficiency range of 0.001 to 0.094, very similar to the state interpreted as acceptor photophysics in Fig. 3. In order to confirm this, we look at the transition rates between state 1 and state 3. From state 3 to state 1, the transition rate has 95% credible interval (0.003 to 0.38) ms^−1^ while the reverse transition has (0.09 to 5.96) ms^−1^. Both are similar to the photophysical rate ranges reported in Fig. 3, and subtle discrepancies may originate from variations in the acceptors’ photophysics as they are brought closer to the donor dyes in the ProT*α*-H1 complex.

On the flip side, interpreting state 2 is trickier as both Figs. 4a and 4d indicate that the system hardly spends any time in state 2 (average lifetime is about 0.01 ms). However, state 2’s parameter posterior probability distributions, Figs. 4d to 4f, despite being broad, do not closely follow the shape of the corresponding priors (for this case see Fig. S.1 state 3), meaning the analyzed bursts probably carry enough data supporting state 2’s existence but its reaction-diffusion dynamics as wells as FRET efficiency can hardly be pinpointed due to its short lifetime; see Fig. S.4 for how short lifetimes affect our analysis. We also speculate that state 2 may originate from the photoblinking of acceptor fluorophores as fluorophores may exhibit photophysics dependent upon variations in ProT*α* and H1 concentrations. Another speculation is that state 2 corresponds to a free ProT*α* molecule and its short lifetime is caused by the abundance of H1 in the surrounding environment. Though this speculation is less likely, as state 2’s escape rate (Fig. 4d blue) is much higher than the association rates (with 10 nM H1) reported in Ref. [41], we mention it nonetheless due to the high uncertainty in state 2’s escape rate.

### 2.4 50 pM labeled ProT*α*, 20 µM H1, and 84 µM unlabeled ProT*α*

In order to further demonstrate the profound data efficiency of Bayes-smRD, we now move to the third date set in which unlabeled ProT*α* is added (50 pM labeled ProT*α* 20 µM H1, and 84 µM unlabeled ProT*α*). Under this set of concentrations, we expect both ProT*α*-H1 and ProT*α*_2_-H1 to be present [41].

Following the argument we made in the previous subsection, we still run a three-state model with the expectation of seeing three states with different FRET efficiencies and subsequently interpret these states based on their reaction-diffusion dynamics. Immediately from Figs. 5a and 5f we can see that, as expected, Bayes-smRD indeed recovers three distinct states whose posteriors over FRET efficiency differ substantially from the prior and captures transitions amongst them. Moreover, Fig. 5d shows that the average lifetimes of the zero FRET (state 1, gray) and the low FRET state (state 2, blue) are both around 0.2 ms. This indicates that, in order to capture these transitions, binning-based FRET analysis tools must have a bin size no greater than 0.2 ms. However, in the burst shown in Fig. 5c, there are 760 photon detections in total and hence setting 0.2 ms means each bin contains less than 40 photons. Such low number of detections will result in very low signal-to-noise ratio thereby greatly undermining the effectiveness of binning-based tools [28] and highlighting the importance of the direct photon-by-photon analysis used herein.

**Figure 5:**
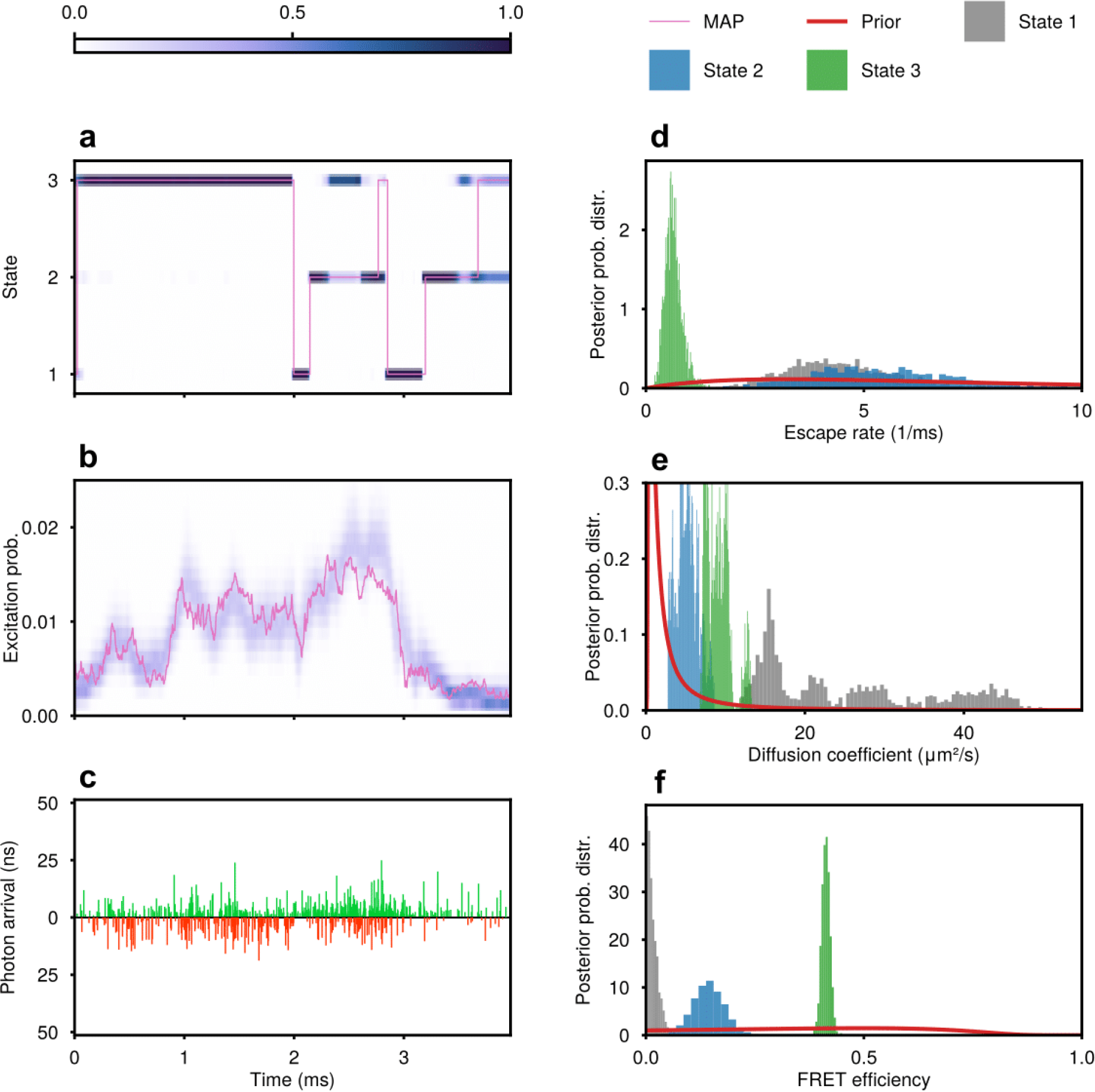
Results from the experimental data set with 50 pM labeled ProT*α* 20 µM H1, and 84 µM unlabeled ProT*α*. Layout is the same as Fig. 3.

In previous subsections, we have presented how the zero-FRET state (Figs. S.1 and 4 state 1) and the low-FRET state (Fig. S.1 state 2) are interpreted as acceptors’ dark state and free ProT*α*, respectively. The same arguments still hold for state 1 and 2 in Fig. 5. Though state 2’s diffusion coefficient and FRET efficiency in this dataset differ from the estimates presented in Figs. S.1 and 3, we attribute this discrepancy to the finiteness of data as only about 7% of the 4548 photons collected in all six bursts originate from state 2. One may argue against this statement by saying the finiteness of data should result in higher uncertainties represented by wider posteriors, unlike what is shown in Fig. 5e. However, a posterior probability distribution’s breadth reflects uncertainty when sufficient samples are present. Therefore, the number of photons and posterior breadths are both necessary in evaluating the confidence of our estimates.

On the other hand, it is still worth discussing what contributes to state 3 in Fig. 5. As mentioned earlier, under this set of concentrations (50 pM labeled ProT*α* 20 µM H1, and 84 µM unlabeled ProT*α*), according to [41], we anticipate complexes including not only ProT*α*-H1 but also larger ternary complexes such as ProT*α*_2_-H1 or ProT*α*-H1_2_ to form (see Fig. 2).

In order to proceed, we first calculate state 3’s escape rate from Fig. 5d, whose 95% credible interval is (0.31 to 1.07) ms^−1^ and mean value is 0.63 ms^−1^. These numbers are smaller than the dissociation rate of ProT*α*_2_-H1 reported in Ref. [41], which is 1.9 ms^−1^. However, we argue this is not a discrepancy. The key point here is that ProT*α*-H1 and ProT*α*_2_-H1 do not show noticeable difference in their FRET efficiencies [41] and hence, they can hardly be separated during burst selection. Therefore, part of the six burst analyzed may contain ProT*α*-H1 while some others contain ProT*α*_2_-H1. Consequently, state 3 in Fig. 5 can represent a mixture of these two complexes. Since ProT*α*-H1 has a much lower dissociation rate, 1.7 ×10^−3^ ms^−1^ [41], it is reasonable that the overall escape rate of state 3 is lower than 1.9 ms^−1^. This explanation in fact highlights the importance of having a tool capable of inferring reaction-diffusion dynamics from single bursts.

### 2.5 Synthetic data

To validate Bayes-smRD, we generate synthetic data by computer simulation [45–49], representing the Brownian motion of a single molecule in a confocal volume while undergoing transitions. In these simulations, we incorporate real-life complications, including background noise, IRF, direct excitation of the acceptor fluorophores, excitation probabilities dependent on light intensity at the fluorophore’s instantaneous spatial location, and detector bleed-through. These parameters mimic those in real experiments [40, 41] and are tabulated in SI table Table S.2.

As illustrated in Fig. 6, Bayes-smRD is able to capture a single molecule’s state and spatial trajectories simultaneously in good agreement with ground truth. In Fig. 6, (d) and (e) show Bayes-smRD’s performance in learning reaction rates (escape rates) as well as diffusion coefficients. All ground truth values fall within the 95% credible intervals, and usually lie close to the posterior probability distribution maxima. Moreover, Fig. 6 shows that Bayes-smRD avoids binning artifacts introduced in FRET analysis methods thereby reducing temporal resolution as transitions occurring on timescales shorter than the bin size are averaged out.

**Figure 6:**
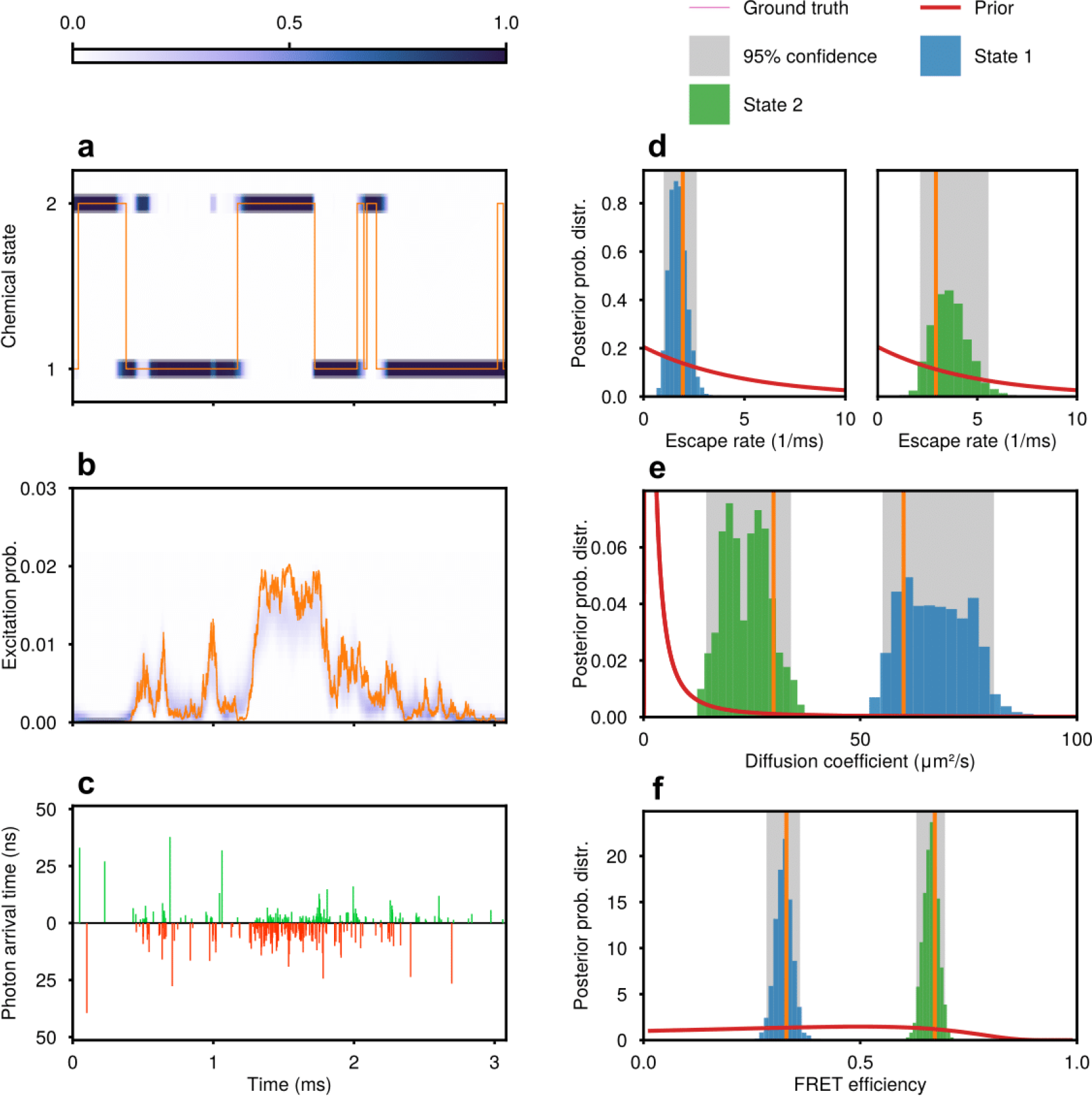
Results from synthetic data, along with the corresponding ground truths, priors, and credible intervals. Our simulation assumes a two-state model. Layout is the same as in Fig. 3.

To further demonstrate Bayes-smRD’s robustness and validity, we tested our method by varying two key quantities, the number of bursts included in the analysis as well as reaction rates. Figures S.3a to S.3c shows how Bayes-smRD performs better (narrower credible intervals centered at ground truths) as more bursts (thus more photons and more information) are considered in the calculation while fixing the excitation rate. In Figs. S.3d to S.3f, we multiply all entries of the original transition rate matrix by a multiplicative factor while keeping the photon counting rate and the total number of photons fixed. This quantifies how higher photon counting rates are needed to probe faster reaction-diffusion dynamics for fixed length trajectories to maintain about the same degree of uncertainty. Roughly, photon counting rates should scale linearly with reaction kinetics.

In addition, we also briefly explain the implication of Figures S.3a to S.3c on diffusion coefficients and excitation rate. As these quantities directly affect photon numbers obtained in each burst, the effect of having a rapidly diffusing species is then equivalent to reducing the photon budget available for analysis within a burst.

Along these lines, we further test Bayes-smRD’s robustness by increasing a state’s lifetime while keeping the other state’s lifetime constant in Figs. S.4a to S.4c, and by increasing the fraction of a burst analyzed in Figs. S.4d to S.4f. The results from both tests are consistent with our conclusion, that is photon number and photon counting rates both positively contribute to our analyses.

## 3 Discussion

The current focus in unraveling single-molecule reaction-diffusion dynamics at the highest temporal resolution necessarily requires the analysis of sequential photon arrivals. Single-photon analysis, already common in FRET [5, 16–18], is key toward resolving fast processes including protein folding [4] as well as interactions of transcription factors with nucleotides critical to cellular regulation [50]. On the other hand, a single-molecule data analysis method capable of learning reaction-diffusion dynamics can be coupled with deep learning based tools predicting biochemical reactions [51] to provide training and verification data. While many single-molecule processes can be probed by FRET-labeled molecules tagged to surfaces [25, 26, 41] in addition to force spectroscopy [52], or both combined [53, 54], these setups also raise questions as to whether fixing molecules to surfaces impact their reactivity [30].

Diffusion-limited single-molecule reactions, with both molecular actors freely diffusing, however introduce unique complexities that so far have limited our ability to resolve smRD models from photon arrivals.

Currently, in an effort to achieve smRD, severe approximations must be invoked including assuming that the illumination is roughly uniform over the diffusion volume [5, 16], thereby eliminating our ability to infer spatial trajectories containing information on state switching.

Alternatively, tens of millions of photons have to be used in the analysis thereby limiting our ability to resolve events at the single-molecule level [4, 41]. At its very core is the problem that a molecule diffuses in continuous space across an inhomogeneously illuminated volume while evolving in a discrete state space.

As we work within the Bayesian paradigm, Bayes-smRD can rigorously propagate all sources of error including spectral bleed-through and background photons into estimates of the reaction rates, diffusion coefficients, FRET efficiencies of each state. Inevitably, Bayes-smRD comes with higher computational cost as compared with traditional correlative methods [55, 56]. Bayes-smRD’s computational time scales roughly linearly with the number of pulses considered.

Beyond providing a single-molecule reaction-diffusion picture, Bayes-smRD can be further generalized to include any confocal volume shape by inserting its mathematical form into Eqs. (S.26) and (S.27). What is more, the method can also treat any pulse shape or sequence by appropriate modification of Eqs. (S.44) to (S.46).

Despite advancements brought by Bayes-smRD, several challenges remain (beyond reactions occurring on experimentally inaccessible timescales): (i) finite data; (ii) multiple labeled particles present in the observation volume; and (iii) unknown number of system states.

Challenge (i) may arise for different reasons, including but not limited to, fast diffusion dynamics regarding the size of observation volumes and rapid photobleaching. In this paper, in order to ease the issue, we choose to analyze several bursts together assuming all selected bursts share the same reaction-diffusion dynamics. An alternative would be to rely on increasingly photostable labels [57] as well as detectors with shorter dead times, to help maximize the information contained in each burst, though theoretical tools may be leveraged to partially beat detector dead times or even integration times in the case of binned photon analysis [58, 59].

On the other hand, challenge (ii) and (iii) can potentially be resolved theoretically. In the Bayesian paradigm, particle number and state number can both be treated as parameters to learn. This approach, requiring Bayesian nonparametrics, would place priors on infinite candidate number of states and molecules, *i*.*e*., it would require a doubly nonparametric framework. Despite technical challenges, the ability to theoretically recruit, within a computational framework, multiple states would provide a powerful generalization.

While Bayes-smRD has been applied here to protein binding interactions by monitoring labels from just one binding partner, it raises broader questions. In particular, it places within conceptual reach the notion that dynamics may be extracted from multiple molecular actors all simultaneously operating within a small diffraction limited region, such as a gene locus under active transcription [60], from photon arrivals.

## 4 Methods

### 4.1 Mathematical formulation

We immediately consider equally spaced pulses at times *t*_1:*K*_ = (*t*_1_, *t*_2_, …, *t*_*K*_) with an interpulse interval of *T* and total pulse number *K*. At time point *t*_*k*_ we have three observations: (i) if there is a photon detection; (ii) the photon arrival time following the pulsed laser *δ*_*k*_, termed the microtime; and (iii) the detector in which the photon is detected *d*_*k*_ (correlated to the color of the photon). In case pulse *k* does not yield a photon detection, both *δ*_*k*_ and *d*_*k*_ take the value of void (∅). These three types of observations can be combined into two arrays: *δ*_1:*K*_ = (*δ*_1_, *δ*_2_, …, *δ*_*K*_) and *d*_1:*K*_ = (*d*_1_, *d*_2_, …, *d*_*K*_).

Now, both *δ*_1:*K*_ and *d*_1:*K*_ are used as the input data in order to construct a posterior over the quantities we care about, namely, ***θ*** = (**Q**_*c*_, *D*_1:*M*_, *c*_1:*K*_, **x**_1:*K*_). Here, **Q**_*c*_ is the state transition rate matrix of size *M × M*, where *M* is the number of states, *D*_1:*M*_ contains all the diffusion coefficients of each state, *c*_1:*K*_ and **x**_1:*K*_ are the state and spatial trajectories, respectively.

We assume that states and spatial positions remain relatively constant during the interpulse interval of approximately 50 ns. To validata our assumption, we perform two quick back-of-envelope calculations. After transitioning into a state with a lifetime of 5 µs, the probability of the system leaving this state within 50 ns is less than 1%. Furthermore, if a particle in this state has a diffusion coefficient of 1000 µm^2^s^−1^, there is a 95% chance that the particle’s displacement will be less than 0.02 µm, which is more than 10 times smaller than the size of a confocal spot.

#### 4.1.1 Forward Model

In order to simulate and later infer ***θ***, we must first prescribe the system’s evolution, in terms of its state as well as spatial trajectories. The state trajectory is given by

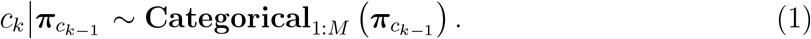

Here, 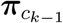 is the row *c*_*k−*1_ of the transition probability matrix **Π**, and **Π** is obtained from **Π** = exp(*T* **Q**_*c*_). Next, the evolution of the spatial trajectory is given by

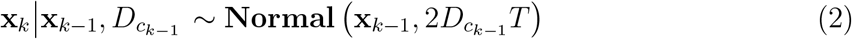

which follows from the solution of the diffusion equation with open boundaries.

With state and spatial trajectories, we can briefly describe how they give rise to observations *δ*_*k*_ and *d*_*k*_. As mentioned earlier, *d*_*k*_ has 3 possible outcomes, no photon detected (*d*_*k*_ = ∅, a photon detected in donor channels (*d*_*k*_ = *D*), or a photon detected in acceptor channels (*d*_*k*_ = *A*), so *d*_*k*_ is sampled from

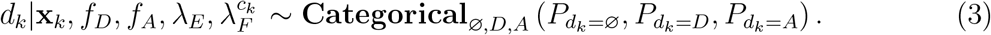

Here, 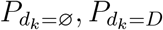 and 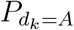 are probabilities of each possible outcome whose complete formulas can be found in Eqs. (S.44) to (S.46), and *f*_*D*_ and *f*_*A*_ are two binary variables marking the existence of donor and acceptor, respectively. These probabilities are calculated based on the state *c*_*k*_, the illumination at position **x**_*k*_, a Bernoulli random variable denoting whether an active donor/acceptor fluorophore is present or not *f*_*D/A*_, the effective excitation rate *λ*_*E*_, and the FRET rates of each state 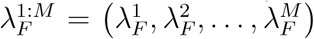 Here, we define the effective excitation rate as the number of molecule-emitted photon detections per u it time.

Also, in our model, the true FRET efficiency of state *k* is defined as 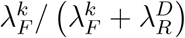 where 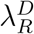 is the predefined donor emission rate.

In addition, calculating these probabilities also involves the consideration of spectral bleed-through, uniform background photons, and the direct excitation of acceptor dyes, see SI Section 2 for details. For bleed-through, we predefine probabilities connecting photon emission channels to detection channels. Uniform background photons are treated as independent photon sources with fixed emission rates. Furthermore, the direct excitation of acceptor dyes is dealt with by introducing a multiplicative factor multiplying the donor excitation rate. All predefined parameters can be obtained based on the wavelengths of photon emissions and optical filters used in actual experiments.

Next we write down a likelihood for the microtime observation *δ*_*k*_. When no photon is detected (*d*_*k*_ = ∅), *δ*_*k*_ is identically void (∅) with probability 1. Otherwise, *δ*_*k*_ is sampled from

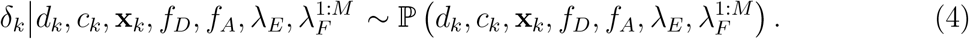

The full expression for this distribution can be found in Eqs. (S.21) to (S.23). Eq. (4) is derived mostly from the convolutions of exponential distributions and the IRF. The exponential distributions reflect waiting times in the excited states of both donor and acceptor, so they depend on 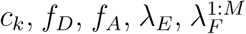 as well as the pre-calibrated donor and acceptor emission rates 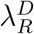 and 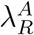 Moreover, the IRF, whose mathematical form is pre-calibrated from experiments, is characterized by two fixed parameters, offset *τ*_*δ*_ and width *σ*_*δ*_.

#### 4.1.2 Inverse Model

From the previous section, we see that in order to infer ***θ***, we must also learn the latent variables coinciding with the state trajectory *c*_1:*K*_ and spatial trajectory **x**_1:*K*_. As such, the full set of quantities to be learned can now be expanded to 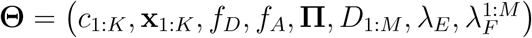 Given all the equations above, and the expanded likelihoods accounting for bleed-through, uniform background photons, acceptor direct excitation, and IRFs, see Eqs. (S.47) to (S.49), we can now, alongside with priors, construct a full posterior probability distribution over ***θ***. In order to sample this high dimensional posterior, *P* (**Θ** *d*_1:*K*_, *δ*_1:*K*_), we opt for specialized Markov Chain Monte Carlo schemes that we design herein. The sampling of this posterior is achieved by invoking a global Gibbs sampling scheme [61]. Within this Gibbs sampler, for *f*_*D*_, *f*_*A*_, each row of **Π**, and each element of *D*_1:*M*_, we select priors listed in Section B.6. Briefly, for computational convenience (detailed in the Supplementary Information), directly sampling *f*_*D*_ and *f*_*A*_ requires Bernoulli priors, each row of **Π** requires a Dirichlet prior, and each element of *D*_1:*M*_ requires an Inverse Gamma prior. The corresponding conditional posteriors can be found in Eqs. (S.52), (S.53), (S.116) and (S.125). As for *c*_1:*K*_, we place a Categorical distribution as prior on *c*_1_, and then apply the forward-filtering backward-sampling algorithm [62] to sample all the states recursively, as shown in Eq. (S.111).

Other quantities, 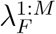, **x**_1:*K*_ and *λ*_*E*_, cannot be sampled directly. As a result, we use the Metropolis-Hastings (MH) algorithm [63, 64]. To sample 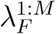, we use Gamma distributions as proposal probability distributions and prior probability distributions. As **x**_1:*K*_ and *λ*_*E*_ are slow to converge, on account of the fact **x**_1:*K*_ contains many intercorrelated continuous variables and inferring *λ*_*E*_ heavily depends on **x**_1:*K*_, we therefore opt for Hamiltonian Monte Carlo (HMC) steps [65] within our broader Gibbs sampling scheme to help speed up the overall convergence. Detailed sampling schemes for 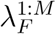, **x**_1:*K*_ and *λ*_*E*_ are covered in Section C.3, Section C.7, and Section C.8.

### 4.2 Burst selection

Burst selection is achieved by first binning the single-photon arrival data and setting a threshold on the number of photons per bin. For bins above the threshold, we then locate a bursts’ start and end times. Within each burst, we calculate the FRET efficiencies and stoichiometries with the corrections often regarding background photon emission rate, detection efficiencies, and bleed-through ratios provided by our experimental collaborators. For the experimental data sets analyzed in this paper, corrections are applied through the route correction matrix [66, 67]. FRET efficiencies and stoichiometries are checked to rule out any burst that may contain uneven numbers of donor fluorophores and acceptor fluorophores.

## 5 Acknowledgements

We all thank Benjamin Schuler, Daniel Nettels, and Andrea Sottini for interesting discussions as well as the data provided. We also further thank Ioannis Sgouralis, Mohamadreza Fazel, Ayush Saurabh, Shep Bryan, Camille Moyer, Matthew Safar, and Max Schweiger for helpful discussions. SP acknowledges support from the NIH (grant no. R01GM134426, R01GM130745, and R35GM148237).

## 6 Code availability

Our code is available at https://github.com/LabPresse/Bayes-smRD.

## 7 Data availability

Synthetic datasets are generated with the code provided in the previous section, experimental datasets come from Ref. [41].

## 8 Author contributions

S.P. conceived and supervised the project, provided all the computational resources, and revised the manuscript. S.J. and Z.K. provided codes from their previous works, co-supervised the project, and revised the manuscript. L.X. wrote the code, performed data analyses, and wrote the manuscript.

