## Supplementary Information for "Single-Molecule Reaction-Diffusion"

### 1 Supplementary figures

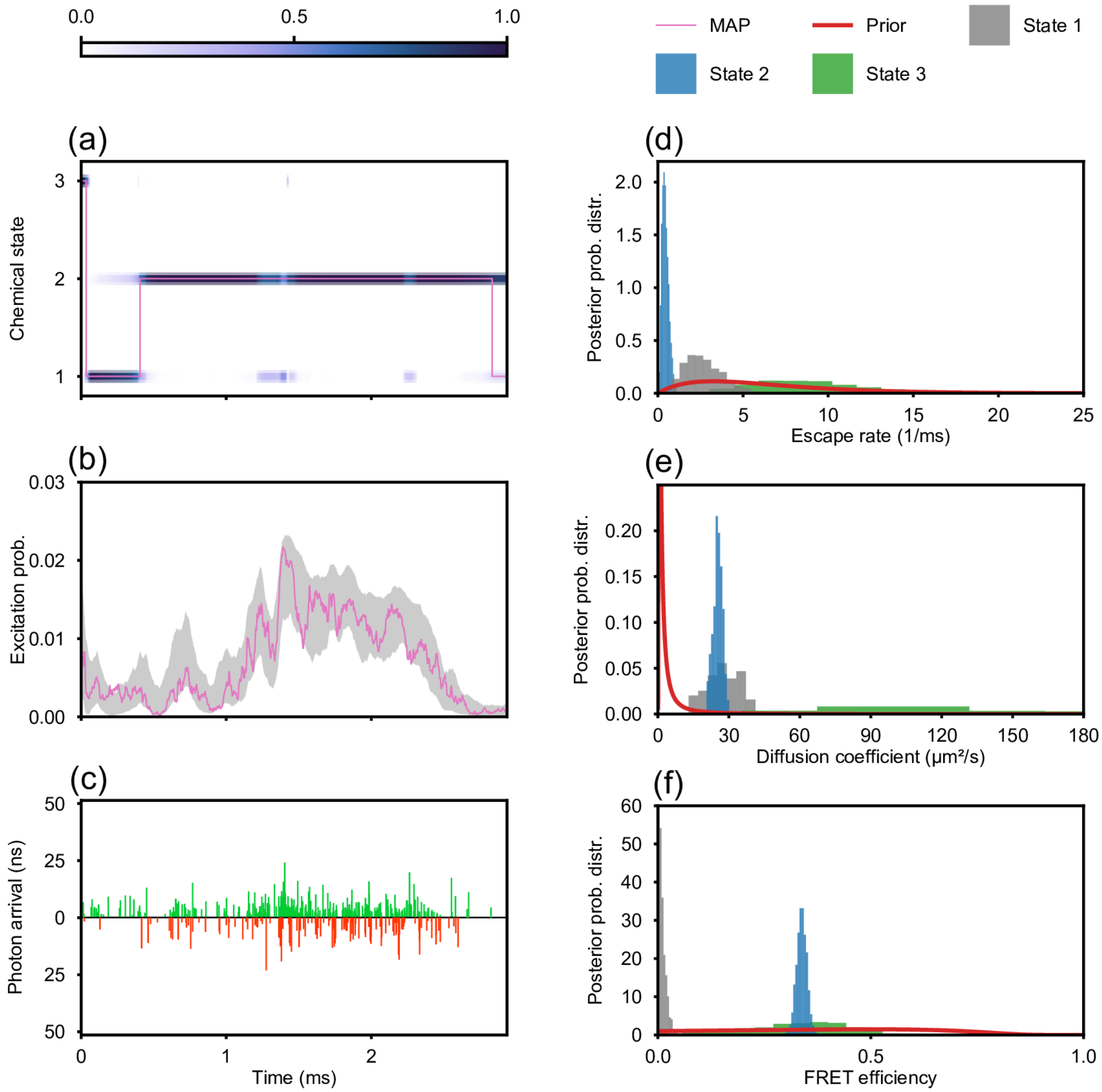


Figure S.1: Results from the experimental data set with 75 pM P* only. Layout is the same as in Fig. 3.

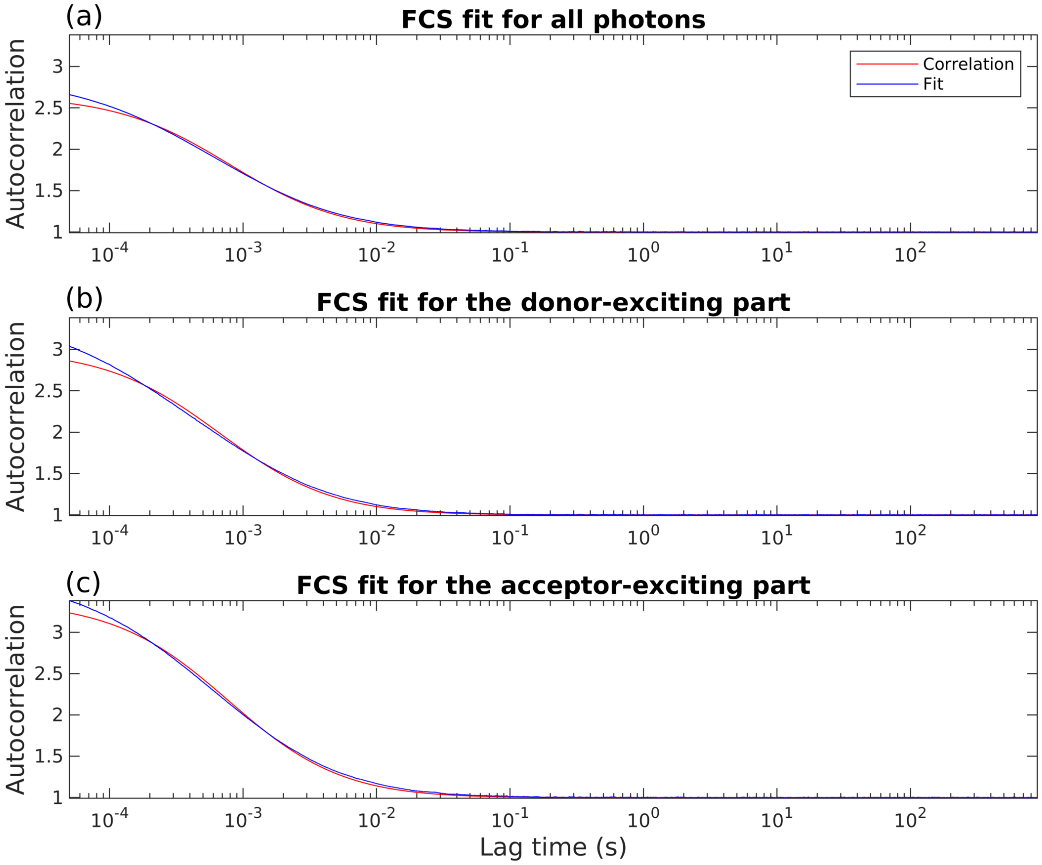


Figure S.2: FCS analysis of the data set analyzed in Fig. 3.

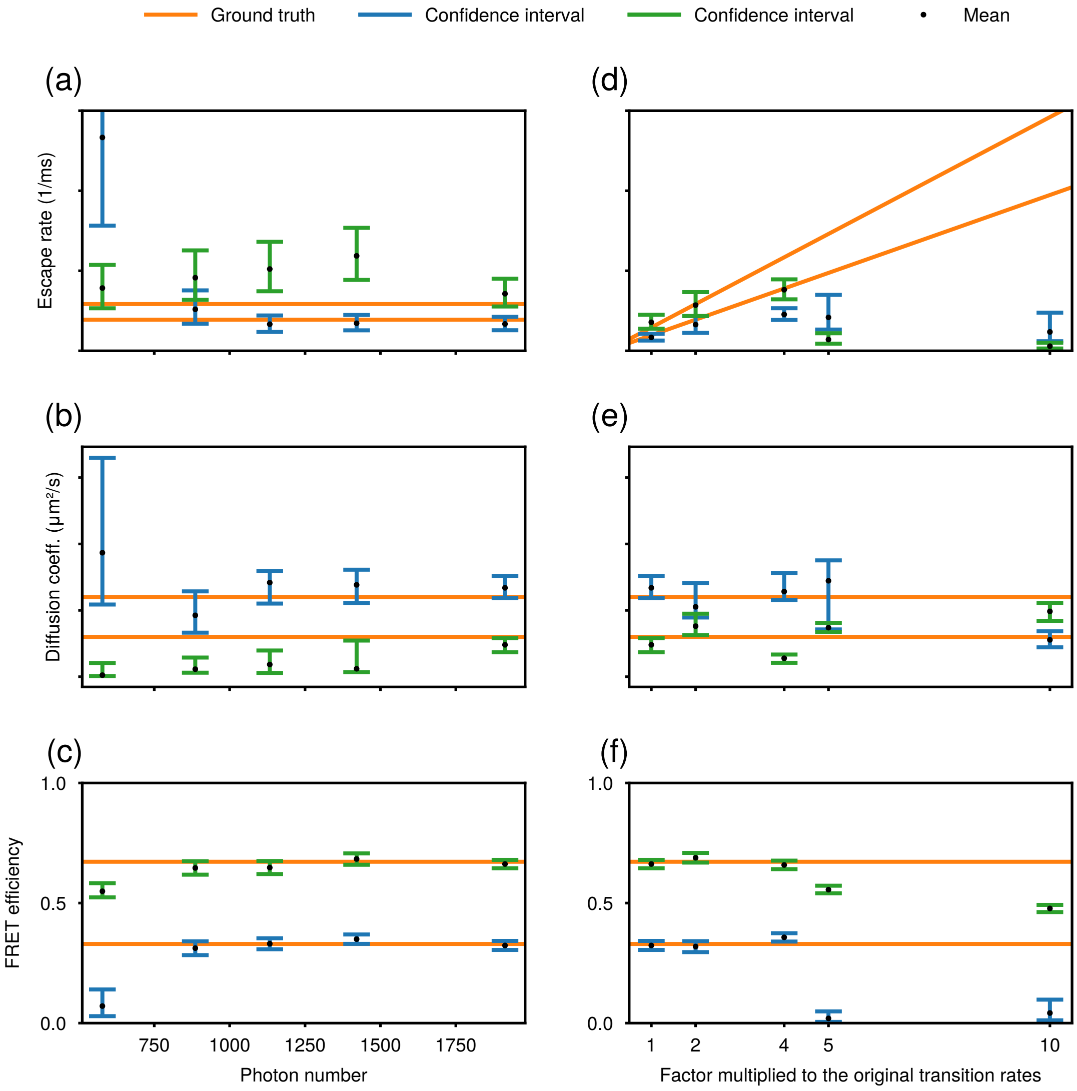


Figure S.3: Tests of our method's robustness. (a) Our method works better as more bursts (thus more photons and naturally more information) are included in the analysis. (b) Here, we gradually increase all transition rates by a factor while holding the photon counting rate fixed. Our method starts to fail when both transitions rates approach the reciprocal of the average inter-photon arrival time.

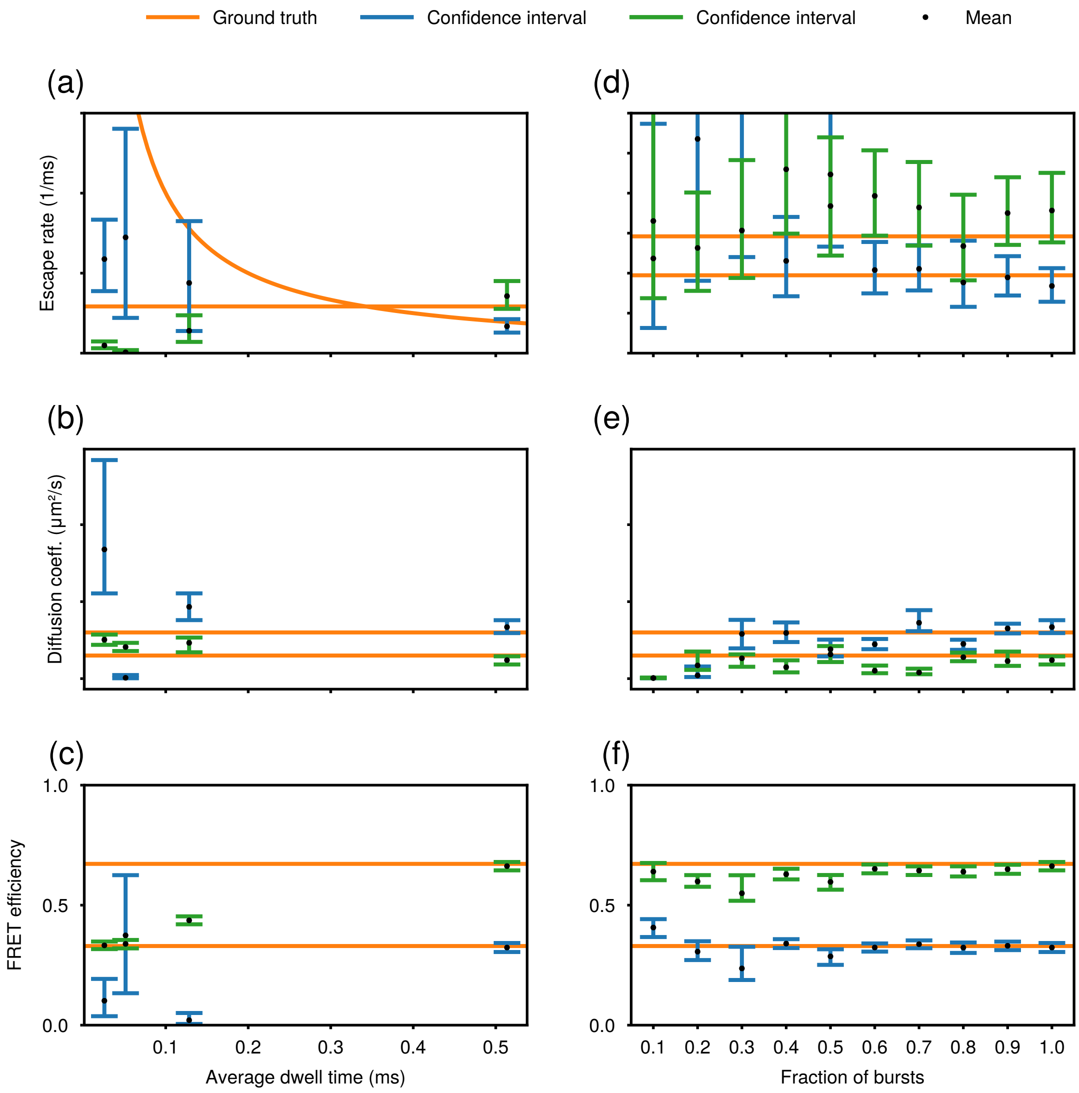


Figure S.4: More tests of our method's robustness. (a) Here, we gradually increase the average dwell time (lifetime) of one state. Our method starts to capture correct reaction-diffusion dynamics the average lifetime becomes longer than the inter-photon arrival times. (b) Our method works better longer bursts (thus more photons considered and more parts of confocal volume explored) are included in the analysis.

### 2 Synthetic data generation

#### B.1 Transition probability matrix

We start by defining the total number of states $M$ for the molecule of interest along with the corresponding $M \times M$ state transition rate matrix $\mathbf{Q}_c = (\lambda_{i,j})$. Here, $\lambda_{i,j}$ represents the instantaneous transition rate from state $i$ to state $j$ if $i \neq j$, otherwise, $\lambda_{i,i} = -\sum_{m=1,m\neq i} \lambda_{i,m}$. For a molecule with rates independent of time over the course of an experiment, we can calculate the state transition probability matrix between two successive pulses $\mathbf{\Pi}$ by

$$\mathbf{\Pi} = \exp\left(T\mathbf{Q}_c\right) \tag{S.1}$$

where $T$ is the interpulse interval, or the pulse period. An element of $\mathbf{\Pi}$, say $\pi_{i,j}$, stands for the probability of the molecule being in state $j$ at the next pulse given its state being $i$ at the previous one.

#### B.2 State trajectory

Next, we can use $\mathbf{\Pi}$ to produce the molecule's state at each pulse. As such, we start with the definition of the state trajectory $c_{1:K}$ and define $c_k$ as the molecule's state at the $k$-th pulse. In order to generate the full trajectory, we first sample $c_1$ from the discrete uniform distribution over $1, 2, \ldots, M$, then, all the subsequent states are sampled from

$$c_k \big| \boldsymbol{\pi}_{c_{k-1}} \sim \mathbf{Categorical}_{1:M}\left(\boldsymbol{\pi}_{c_{k-1}}\right) \tag{S.2}$$

for $k = 2, 3 \ldots, K$. Here $\boldsymbol{\pi}_{c_{k-1}}$ is row $c_{k-1}$ of $\mathbf{\Pi}$.

#### B.3 Spatial trajectory

The molecule's spatial trajectory is generated recursively in a similar fashion. We set up the coordinate system such that the center of the confocal volume is the origin. (Here, we define the origin as the spot with the maximum illumination.) The initial position is sampled from three Normal distributions with zero mean, and $\sigma_x^2$, $\sigma_y^2$, $\sigma_z^2$ as variances,

$$\begin{aligned} x_1 &\sim \mathbf{Normal}\left(0, \sigma_x^2\right), \\ y_1 &\sim \mathbf{Normal}\left(0, \sigma_y^2\right), \\ z_1 &\sim \mathbf{Normal}\left(0, \sigma_z^2\right). \end{aligned} \tag{S.3}$$

According to Fick's law of diffusion, and with the predefined diffusion coefficients $D_{1:M}$, we sample the rest of the trajectory from

$$\mathbf{x}_k \big| \mathbf{x}_{k-1}, D_{c_{k-1}} \sim \mathbf{Normal}\left(\mathbf{x}_{k-1}, 2D_{c_{k-1}}T\right). \tag{S.4}$$

#### B.4 Photon detection channel and arrival time

At any given time $t$ and position $\mathbf{x}$, provided the system's point spread function (PSF), the instantaneous excitation rate of a donor fluorophore is $f_D\lambda_{E,0}\mathrm{PSF}\left(\mathbf{x}, t\right)$, where $f_D$ is a Bernoulli random variable denoting whether an active donor is present and $\lambda_{E,0}$ is the dye's excitation rate at the center of confocal volume. Then after a very short exposure period, $\mathrm{d}t$, the dye's probability of staying within its ground state reads $1 - \exp\left[-f_D\lambda_{E,0}\mathrm{PSF}\left(\mathbf{x}, t\right)\mathrm{d}t\right]$.

In practice, for pulsed illumination, the duration of a pulse is typically on the order of picoseconds, which is too short for a molecule to move substantially and to emit. Thus, the probability of a donor staying in its ground state during the $k$-th pulse can be calculated as $1-\exp\left[-f_D\lambda_{E,0}\tau\text{PSF}\left(\mathbf{x}\right)\right]$. Here, we also approximate the pulses as square waves with width, termed pulse width, $\tau$. In reality, only a fraction ($\phi$) of excitations lead to photon detections. In order to incorporate this effect, we define the effective excitation rate $\lambda_E$ such that

$$\phi\left[1-e^{-\lambda_{E,0}\tau\text{PSF}(\mathbf{x})}\right]=1-e^{-\lambda_E\tau\text{PSF}(\mathbf{x})}. \tag{S.5}$$

That is to say, $\lambda_E$ is the rate of detected donor excitation at the center of the PSF, and $1-\exp\left[-f_D\lambda_E\tau\text{PSF}\left(\mathbf{x}\right)\right]$ is the probability of getting nonzero photon detections originating from direct donor excitations after a single pulse. The calculation we've done so far is also valid for an acceptor fluorophore, but instead of introducing another effective excitation rate for the acceptor, we simply assume the rate is proportional to $\lambda_E$ with a pre-calibrated multiplicative factor $\xi$. Consequently, $1-\exp\left[-f_A\xi\lambda_E\tau\text{PSF}\left(\mathbf{x}\right)\right]$ is the probability of getting nonzero photon detections originating from direct acceptor excitations after a single pulse.

Besides direct donor excitations and direct acceptor excitations, a detected photon may also come from the uniform donor channel background and the uniform acceptor channel background. By definition, probabilities of getting nonzero photon detections from them within in one interpulse interval are $1-\exp\left(-\lambda_B^D T\right)$ and $1-\exp\left(-\lambda_B^A T\right)$. Here, $\lambda_B^D$ and $\lambda_B^A$ are the effective donor background emission rates and the effective acceptor background emission rates, respectively.

For notational simplicity alone, we label the aforementioned photon origins as: $o=1$ (direct donor excitation); $o=2$ (direct acceptor excitation); $o=3$ (uniform donor channel background); and $o=4$ (uniform acceptor channel background). As the probability of detecting a photon is no greater than 0.02 in the experimental data sets analyzed in this manuscript (Figs. 3, 4, and 5), we omit the case of getting multiple photon detections within one interpulse interval. Furthermore, the dead time of single-photon detectors also prevents the detection of multiple photons if they are temporally close. We therefore assume, in the interpulse interval following the $k$-th pulse, the corresponding origin $o_k$ can either be a number between 1 and 4 or be $\varnothing$ which represents no photon detection.

In order to move forward in generating actual observations, we still need to write down a probability distribution over each $o_k$. Here, we make another assumption that fluorophore fluorescence lifetimes are so small compared to the interpulse interval that all the dyes will always be in their ground states by the occurrence of the next pulse. This is assumption is valid for fluorophores such as Alexa 488/594 (used in the experimental data set) have lifetimes shorter than 5 ns [1], and hence the chance for them to stay in excited states for an entire interpulse interval (about 50 ns) is less than $10^{-4}$. Under this assumption, $o_k$ only depends on the current state of the system, and hence, if a molecule carries both a donor and an acceptor, we can sample $o_k$ from

$$o_k|\mathbf{x}_k, f_D, f_A, \lambda_E \sim \textbf{Categorical}_{\varnothing,1:4}\left(P_{o_k=\varnothing}, P_{o_k=1}, P_{o_k=2}, P_{o_k=3}, P_{o_k=4}\right) \tag{S.6}$$

where

$$P_{o_k=\varnothing} = \left(e^{f_D \lambda_E \tau \mathrm{PSF}_k} + e^{f_A \xi \lambda_E \tau \mathrm{PSF}_k} + e^{\lambda_B^D T} + e^{\lambda_B^A T} - 3\right)^{-1} \tag{S.7}$$

$$P_{o_k=1} = \left(e^{f_D \lambda_E \tau \mathrm{PSF}_k} - 1\right) P_{o_k=\varnothing} \tag{S.8}$$

$$P_{o_k=2} = \left(e^{f_A \xi \lambda_E \tau \mathrm{PSF}_k} - 1\right) P_{o_k=\varnothing} \tag{S.9}$$

$$P_{o_k=3} = \left(e^{\lambda_B^D T} - 1\right) P_{o_k=\varnothing} \tag{S.10}$$

$$P_{o_k=4} = \left(e^{\lambda_B^A T} - 1\right) P_{o_k=\varnothing} \tag{S.11}$$

Subsequently, we generate the output for photon detection channels, $d_k$, and arrival times, $\delta_k$, for each case of $o_k$. Here, allowed values for each $d_k$ include no detection ($\varnothing$), the donor channel ($D$), and the acceptor channel ($A$); and $o_k$ can either be $\varnothing$ or a real number. Here $o_k = \varnothing$ corresponds to no photon detection, so

$$P\left(d_k | o_k = \varnothing\right) = \delta_{d_k,\varnothing}, \tag{S.12}$$

$$P\left(\delta_k | o_k = \varnothing\right) = \delta_{\delta_k,\varnothing}, \tag{S.13}$$

where $\delta_{x,y}$ is the Kronecker delta function. For the uniform background photon ($o_k = 3, 4$) cases, by definition, we have

$$P\left(d_k | o_k = 3\right) = \delta_{d_k,D}, \tag{S.14}$$

$$P\left(d_k | o_k = 4\right) = \delta_{d_k,A}, \tag{S.15}$$

$$P\left(\delta_k | o_k = 3, 4\right) = \frac{1}{T}. \tag{S.16}$$

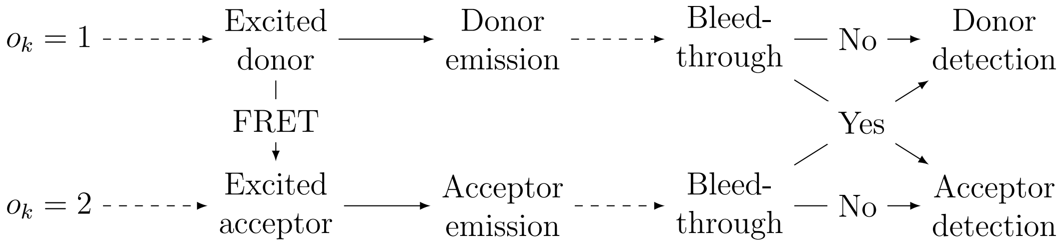


Figure S.5: Different pathways of photon detection from $o_k = 1, 2$. Each solid arrow stands for a process contributing to photon arrival times. Wherever there are two pathways (arrows) coming out from the same source, we need to sample from a Bernoulli distribution to decide which pathway to follow. The arrows between bleed-through and detection come from the IRF.

It is more complicated when a photon detection originates from the fluorophores on a molecule ($o_k = 1, 2$). Fig. S.5 lists all pathways from these origins to different detection channels and is helpful in allowing us to write down the desired probability distributions. First, we specify the Bernoulli distributions involved in Fig. S.5. Given $o_k = 1$ and a state $c_k$, whether FRET occurs or not is sampled from a Bernoulli distribution, with

$$P\left(\text{FRET occurs}|o_k = 1, c_k\right) = \frac{f_A \lambda_F^{c_k}}{f_A \lambda_F^{c_k} + \lambda_R^D}. \tag{S.17}$$

The other Bernoulli distributions are tied to detector bleed-through. Here, we define 4 probabilities, $P_{DD}$, $P_{DA}$, $P_{AD}$, and $P_{AA}$, where $P_{XY}$ represents the probability of a photon of channel $X$ being detected as in channel $Y$. Following all pathways in Fig. S.5, we have

$$P\left(d_k|o_k = 1\right) = \frac{1}{f_A \lambda_F^{c_k} + \lambda_R^D}\left(\lambda_R^D P_{DD} + f_A \lambda_F^{c_k} P_{AD}\right)^{\delta_{d_k,D}}\left(\lambda_R^D P_{DA} + f_A \lambda_F^{c_k} P_{AA}\right)^{\delta_{d_k,A}}, \tag{S.18}$$

$$P\left(d_k|o_k = 2\right) = P_{AD}^{\delta_{d_k,D}} P_{AA}^{\delta_{d_k,A}}. \tag{S.19}$$

Fluorescence lifetime probability distributions are also calculated according to these pathways. For instance, the pathway with $o_k = 1$, no FRET, and $d_k = D$ contains donor emission time and IRF. The former follows the exponential distribution with $\lambda_R^D + f_A \lambda_F^{c_k}$ as its rate, and the IRF is a Gaussian whose mean $\tau_\delta$ and standard deviation $\sigma_\delta$ are measured separately. Taking the convolution of them yields the exponentially modified Normal distribution, $g\left(\delta_k; \lambda_R^D + f_A \lambda_F^{c_k}, \tau_\delta, \sigma_\delta\right)$ that is defined as

$$g\left(\delta; \lambda, \tau, \sigma\right) = \frac{\lambda}{2} e^{-\lambda\left(\delta - \tau - \lambda\sigma^2/2\right)} \text{erfc}\left(\frac{-\delta + \tau + \lambda\sigma^2}{\sqrt{2\sigma^2}}\right). \tag{S.20}$$

And the probability of neither FRET nor donor-to-acceptor bleed-through occurring is $\frac{\lambda_R^D}{\lambda_R^D + f_A \lambda_F^{c_k}} \times P_{DD}$.

The other possible way to get $d_k = D$ with $o_k = 1$ is to have both FRET and acceptor-to-donor bleed-through occur. The total arrival time then comes from the convolution of the holding time before FRET, the acceptor lifetime, and IRF, yielding

$$\frac{1}{\lambda_R^D + f_A\left(\lambda_F^{c_k} - \lambda_R^A\right)}\left[g\left(\delta_k; \lambda_R^D + f_A \lambda_F^{c_k}, \tau_\delta, \sigma_\delta\right) - g\left(\delta_k; f_A \lambda_R^A, \tau_\delta, \sigma_\delta\right)\right],$$

with probability $\frac{f_A \lambda_F^{c_k}}{\lambda_R^D + f_A \lambda_F^{c_k}} \times P_{AD}$. Note that a special case occurs when $\lambda_R^D + f_A\left(\lambda_F^{c_k} - \lambda_R^A\right) = 0$, and the distribution takes a different form. As this case almost never happens, we do not provide its derivation.

To summarize, the probability distribution of single-photon arrival time given that the donor is excited and a donor photon is detected is

$$
\begin{aligned}
&P\left(\delta_k|d_k=D,o_k=1,f_A,\lambda_F^{c_k}\right)\\
&=\frac{\lambda_R^D P_{DD}}{\lambda_R^D P_{DD}+f_A\lambda_F^{c_k}P_{AD}}g\left(\delta_k;\lambda_R^D+f_A\lambda_F^{c_k},\tau_\delta,\sigma_\delta\right)\\
&-\frac{f_A\lambda_F^{c_k}P_{AD}}{\lambda_R^D P_{DD}+f_A\lambda_F^{c_k}P_{AD}}\frac{f_A\lambda_R^A}{\lambda_R^D+f_A\left(\lambda_F^{c_k}-\lambda_R^A\right)}g\left(\delta_k;\lambda_R^D+f_A\lambda_F^{c_k},\tau_\delta,\sigma_\delta\right)\\
&+\frac{f_A\lambda_F^{c_k}P_{AD}}{\lambda_R^D P_{DD}+f_A\lambda_F^{c_k}P_{AD}}\frac{\lambda_R^D+f_A\lambda_F^{c_k}}{\lambda_R^D+f_A\left(\lambda_F^{c_k}-\lambda_R^A\right)}g\left(\delta_k;f_A\lambda_R^A,\tau_\delta,\sigma_\delta\right). \qquad \text{(S.21)}
\end{aligned}
$$

Similarly, the probability distribution of single-photon arrival time given that the donor is excited and an acceptor photon is detected is

$$
\begin{aligned}
&P\left(\delta_k|d_k=A,o_k=1,f_A,\lambda_F^{c_k}\right)\\
&=\frac{\lambda_R^D P_{DA}}{\lambda_R^D P_{DA}+f_A\lambda_F^{c_k}P_{AA}}g\left(\delta_k;\lambda_R^D+f_A\lambda_F^{c_k},\tau_\delta,\sigma_\delta\right)\\
&-\frac{f_A\lambda_F^{c_k}P_{AA}}{\lambda_R^D P_{DA}+f_A\lambda_F^{c_k}P_{AA}}\frac{f_A\lambda_R^A}{\lambda_R^D+f_A\left(\lambda_F^{c_k}-\lambda_R^A\right)}g\left(\delta_k;\lambda_R^D+f_A\lambda_F^{c_k},\tau_\delta,\sigma_\delta\right)\\
&+\frac{f_A\lambda_F^{c_k}P_{AA}}{\lambda_R^D P_{DA}+f_A\lambda_F^{c_k}P_{AA}}\frac{\lambda_R^D+f_A\lambda_F^{c_k}}{\lambda_R^D+f_A\left(\lambda_F^{c_k}-\lambda_R^A\right)}g\left(\delta_k;f_A\lambda_R^A,\tau_\delta,\sigma_\delta\right). \qquad \text{(S.22)}
\end{aligned}
$$

As for the cases where $o_k=2$,

$$
P\left(\delta_k|o_k=2\right)=g\left(\delta_k;\lambda_R^A,\tau_\delta,\sigma_\delta\right). \qquad \text{(S.23)}
$$

#### B.5 Simplifications

So for, we have derived the photon detection channel likelihoods (Eqs. (S.18) and (S.19)) and the single-photon arrival time likelihoods (Eqs. (S.21) to (S.23)) in steps for the sake of intuitiveness. However, it is computationally more convenient to combine them into joint probability distributions, $P\left(d_k,\delta_k|o_k,\mathbf{x}_k,f_D,f_A,\lambda_E,\lambda_F^{c_k}\right)$. This is done according to the Bayes' theorem by combining the aforementioned equations in the form of

$$
P\left(d_k,\delta_k|o_k,\mathbf{x}_k,f_D,f_A,\lambda_E,\lambda_F^{c_k}\right)=P\left(\delta_k|d_k,o_k,\mathbf{x}_k,f_D,f_A,\lambda_E,\lambda_F^{c_k}\right)P\left(d_k|o_k,\mathbf{x}_k,f_D,f_A,\lambda_E\right). \qquad \text{(S.24)}
$$

For instance, $P\left(d_k,\delta_k|o_k=1,\mathbf{x}_k,f_D,f_A,\lambda_E,\lambda_F^{c_k}\right)$ is just the product of Eqs. (S.18) and (S.21).

Another simplification we can perform is the marginalization over the photon origins $o_k$, as they carry little physical significance and are mostly introduced to simplify calculation. We do so by

$$
P\left(d_k,\delta_k|\mathbf{x}_k,f_D,f_A,\lambda_E,\lambda_F^{c_k}\right)=\sum_{o_k}P\left(d_k,\delta_k|o_k,\mathbf{x}_k,f_D,f_A,\lambda_E,\lambda_F^{c_k}\right)P\left(o_k|\mathbf{x}_k,f_D,f_A,\lambda_E,\lambda_F^{c_k}\right) \qquad \text{(S.25)}
$$

where the sum is over all possible values of $o_k$.

Therefore, we have worked out the joint observation likelihoods independent of $o_k$. All of these observation likelihoods are worked out and listed in Section B.6.

#### B.6 Model summary

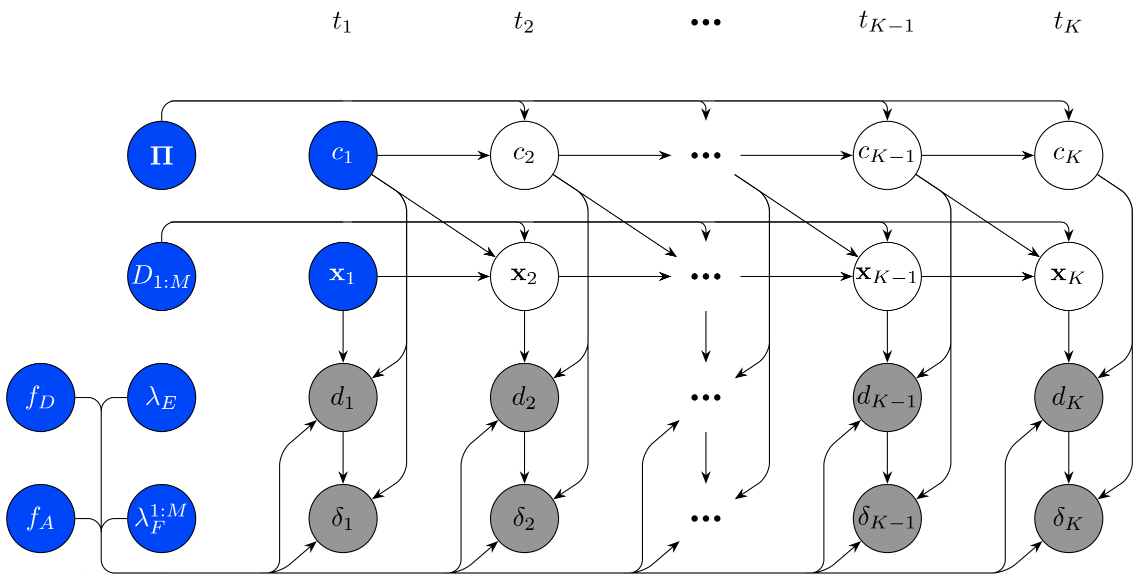


Figure S.6: Bayesian formulation used for the analysis of single-photon arrival data. A single molecule evolves over the experimental time course which is indexed by $k = 1, 2, \dots, K$. Here, $c_k$ and $\mathbf{x}_k$ indicate the state and spatial position of molecule at the $k$-th pulse. During the experiment, two observations (color of detection and single-photon arrival time) $d_k$ and $\delta_k$ are recorded. The state transition probability matrix $\mathbf{\Pi}$ governs the evolution of states while the diffusion coefficients $D_{1:M}$ determine the evolution of the molecular positions. In the graphical model, the measured data are highlighted by gray shaded circles and the model variables, which require priors, are designate by blue circles.

$$\mu_{E,k}^{D} = f_D \lambda_E \mathrm{PSF}_k \tag{S.26}$$

$$\mu_{E,k}^{A} = f_A \xi \lambda_E \mathrm{PSF}_k \tag{S.27}$$

$$\mu_{F}^{c_k} = f_D f_A \lambda_F^{c_k} \tag{S.28}$$

$$\mu_{R}^{D} = f_D \lambda_R^{D} \tag{S.29}$$

$$\mu_{R}^{A} = f_A \lambda_R^{A} \tag{S.30}$$

$$c_1 \sim \mathbf{Categorical}_{1:M}\left(\boldsymbol{\alpha}_c\right) \tag{S.31}$$

$$\boldsymbol{\pi}_m \sim \mathbf{Dirichlet}\left(\boldsymbol{\alpha}_{\boldsymbol{\pi}_m}\right) \tag{S.32}$$

$$c_k \big| \boldsymbol{\pi}_{c_{k-1}} \sim \mathbf{Categorical}_{1:M}\left(\boldsymbol{\pi}_{c_{k-1}}\right) \tag{S.33}$$

$$x_1 \sim \mathbf{Normal}\left(0, \sigma_x^2\right) \tag{S.34}$$

$$y_1 \sim \mathbf{Normal}\left(0, \sigma_y^2\right) \tag{S.35}$$

$$z_1 \sim \mathbf{Normal}\left(0, \sigma_z^2\right) \tag{S.36}$$

$$D_m \sim \mathbf{Inv\text{-}Gamma}\left(\alpha_{D_m}, \beta_{D_m}\right) \tag{S.37}$$

$$\mathbf{x}_k \big| \mathbf{x}_{k-1}, D_{c_{k-1}} \sim \mathbf{Normal}\left(\mathbf{x}_{k-1}, 2D_{c_{k-1}}T\right) \tag{S.38}$$

$$f_D \sim \mathbf{Bernoulli}\left(P_{f_D}\right) \tag{S.39}$$

$$f_A \sim \mathbf{Bernoulli}\left(P_{f_A}\right) \tag{S.40}$$

$$\lambda_E \sim \mathbf{Gamma}\left(\alpha_E, \beta_E/\alpha_E\right) \tag{S.41}$$

$$\lambda_F^m \sim \mathbf{Gamma}\left(\alpha_F^m, \beta_F^m/\alpha_F^m\right) \tag{S.42}$$

$$d_k | \mathbf{x}_k, f_D, f_A, \lambda_E, \lambda_F^{c_k} \sim \mathbf{Categorical}_{\varnothing,D,A}\left(P_{d_k=\varnothing}, P_{d_k=D}, P_{d_k=A}\right) \tag{S.43}$$

$$P_{d_k=\varnothing} = \left(e^{\mu_{E,k}^{D}\tau} + e^{\mu_{E,k}^{A}\tau} + e^{\lambda_B^{D}T} + e^{\lambda_B^{A}T} - 3\right)^{-1} \tag{S.44}$$

$$P_{d_k=D} = P_{d_k=\varnothing}\left[\left(e^{\mu_{E,k}^{D}\tau} - 1\right)\frac{\mu_R^D P_{DD} + \mu_F^{c_k} P_{AD}}{\mu_R^D + \mu_F^{c_k}} + \left(e^{\mu_{E,k}^{A}\tau} - 1\right) P_{AD} + \left(e^{\lambda_B^{D}T} - 1\right)\right] \tag{S.45}$$

$$P_{d_k=A} = P_{d_k=\varnothing}\left[\left(e^{\mu_{E,k}^{D}\tau} - 1\right)\frac{\mu_R^D P_{DA} + \mu_F^{c_k} P_{AA}}{\mu_R^D + \mu_F^{c_k}} + \left(e^{\mu_{E,k}^{A}\tau} - 1\right) P_{AA} + \left(e^{\lambda_B^{A}T} - 1\right)\right] \tag{S.46}$$

$$P\left(\delta_k, d_k=\varnothing|\mathbf{x}_k, f_D, f_A, \lambda_E, \lambda_F^{c_k}\right) = P_{d_k=\varnothing}\delta_{\delta_k,\varnothing}, \tag{S.47}$$

$$\begin{aligned}
&P\left(\delta_k, d_k=D|\mathbf{x}_k, f_D, f_A, \lambda_E, \lambda_F^{c_k}\right)\\
&= P_{d_k=\varnothing}\left(e^{\mu^D_{E,k}\tau}-1\right)\frac{\mu_R^D P_{DD}}{\mu_R^D+\mu_F^{c_k}} g\left(\delta_k;\mu_R^D+\mu_F^{c_k},\tau_\delta,\sigma_\delta\right)\\
&+ P_{d_k=\varnothing}\left(e^{\mu^D_{E,k}\tau}-1\right)\frac{\mu_F^{c_k} P_{AD}}{\mu_R^D+\mu_F^{c_k}}\\
&\times\left[\frac{\mu_R^D+\mu_F^{c_k}}{\mu_R^D+\mu_F^{c_k}-\mu_R^A} g\left(\delta_k;\mu_R^A,\tau_\delta,\sigma_\delta\right) - \frac{\mu_R^A}{\mu_R^D+\mu_F^{c_k}-\mu_R^A} g\left(\delta_k;\mu_R^D+\mu_F^{c_k},\tau_\delta,\sigma_\delta\right)\right]\\
&+ P_{d_k=\varnothing}\left(e^{\mu^A_{E,k}\tau}-1\right)P_{AD} g\left(\delta_k;\mu_R^A,\tau_\delta,\sigma_\delta\right) + P_{d_k=\varnothing}\left(e^{\lambda_B^D T}-1\right)\frac{1}{T},
\end{aligned} \tag{S.48}$$

$$\begin{aligned}
&P\left(\delta_k, d_k=A|\mathbf{x}_k, f_D, f_A, \lambda_E, \lambda_F^{c_k}\right)\\
&= P_{d_k=\varnothing}\left(e^{\mu^D_{E,k}\tau}-1\right)\frac{\mu_R^D P_{DA}}{\mu_R^D+\mu_F^{c_k}} g\left(\delta_k;\mu_R^D+\mu_F^{c_k},\tau_\delta,\sigma_\delta\right)\\
&+ P_{d_k=\varnothing}\left(e^{\mu^D_{E,k}\tau}-1\right)\frac{\mu_F^{c_k} P_{AA}}{\mu_R^D+\mu_F^{c_k}}\\
&\times\left[\frac{\mu_R^D+\mu_F^{c_k}}{\mu_R^D+\mu_F^{c_k}-\mu_R^A} g\left(\delta_k;\mu_R^A,\tau_\delta,\sigma_\delta\right) - \frac{\mu_R^A}{\mu_R^D+\mu_F^{c_k}-\mu_R^A} g\left(\delta_k;\mu_R^D+\mu_F^{c_k},\tau_\delta,\sigma_\delta\right)\right]\\
&+ P_{d_k=\varnothing}\left(e^{\mu^A_{E,k}\tau}-1\right)P_{AA} g\left(\delta_k;\mu_R^A,\tau_\delta,\sigma_\delta\right) + P_{d_k=\varnothing}\left(e^{\lambda_B^A T}-1\right)\frac{1}{T}.
\end{aligned} \tag{S.49}$$

### 3 Detailed inverse model description

#### C.1 The full posterior

The goal of our method, Bayes-smRD, is to sample all the quantities of interest from the full joint posterior probability distribution $P\left(\boldsymbol{\Theta}|d_{1:K},\delta_{1:K}\right)$. Here, the set of all these quantities is denoted as $\boldsymbol{\Theta}$. According to the Bayes' theorem, we have

$$\begin{aligned}
P\left(\boldsymbol{\Theta}|d_{1:K},\delta_{1:K}\right) \propto &\left[\prod_{k=1}^{K} P\left(d_k,\delta_k|\mathbf{x}_k, f_D, f_A, \lambda_E, \lambda_F^{c_k}\right)\right]\left[\prod_{k=2}^{K} P\left(\mathbf{x}_k|\mathbf{x}_{k-1}, D_{c_k}\right) P\left(c_k\middle|\boldsymbol{\pi}_{c_{k-1}}\right)\right]\\
&\times P\left(\mathbf{x}_1\right) P\left(c_1\right) P\left(f_D\right) P\left(f_A\right) P\left(\boldsymbol{\Pi}\right) P\left(\lambda_F^{1:M}\right) P\left(\lambda_E\right) P\left(D_{1:M}\right).
\end{aligned} \tag{S.50}$$

The probability distributions used in Eq. (S.50) have been specified in Section B.6.

As the general form of this posterior is highly complicated, we cannot perform direct sampling. Instead, we proceed with the Gibbs sampling scheme [2] by iteratively updating some quantities of interest from their conditional posteriors. To be more specific, we repeat the following sampling steps:

1. Sample $f_D$ and $f_A$ from $P\left(f_D, f_A\middle|d_{1:K},\delta_{1:K},c_{1:K},\mathbf{x}_{1:K},\boldsymbol{\Pi},D_{1:M},\lambda_E,\lambda_F^{1:M}\right)$;
2. Sample $\mathbf{x}_{1:K}$ from $P\left(\mathbf{x}_{1:K}\middle|d_{1:K},\delta_{1:K},c_{1:K},f_D,f_A,\boldsymbol{\Pi},D_{1:M},\lambda_E,\lambda_F^{1:M}\right)$;

3. Sample $c_{1:K}$ from $P\left(c_{1:K}\middle|d_{1:K}, \delta_{1:K}, \mathbf{x}_{1:K}, f_D, f_A, \mathbf{\Pi}, D_{1:M}, \lambda_E, \lambda_F^{1:M}\right)$;
4. Sample $\mathbf{\Pi}$ from $P\left(\mathbf{\Pi}\middle|d_{1:K}, \delta_{1:K}, c_{1:K}, \mathbf{x}_{1:K}, f_D, f_A, D_{1:M}, \lambda_E, \lambda_F^{1:M}\right)$;
5. Sample $D_{1:M}$ from $P\left(\mathbf{\Pi}\middle|d_{1:K}, \delta_{1:K}, c_{1:K}, \mathbf{x}_{1:K}, f_D, f_A, \mathbf{\Pi}, \lambda_E, \lambda_F^{1:M}\right)$;
6. Sample $\lambda_F^{1:M}$ from $P\left(\mathbf{\Pi}|d_{1:K}, \delta_{1:K}, c_{1:K}, \mathbf{x}_{1:K}, f_D, f_A, \mathbf{\Pi}, D_{1:M}, \lambda_E\right)$;
7. Sample $\lambda_E$ from $P\left(\mathbf{\Pi}\middle|d_{1:K}, \delta_{1:K}, c_{1:K}, \mathbf{x}_{1:K}, f_D, f_A, \mathbf{\Pi}, D_{1:M}, \lambda_F^{1:M}\right)$.

In the following sections we will discuss each step in detail.

#### C.2 Learning fluorophore presence

To construct the marginal posterior in step 1, we simply collect all factors in Eq. (S.50) with $f_D$ or $f_A$ dependence,

$$\begin{aligned} &P\left(f_D, f_A\middle|d_{1:K}, \delta_{1:K}, c_{1:K}, \mathbf{x}_{1:K}, \mathbf{\Pi}, D_{1:M}, \lambda_E, \lambda_F^{1:M}\right) \\ &\propto \left[\prod_{k=1}^{K} P\left(d_k, \delta_k\middle|c_k, \mathbf{x}_k, f_D, f_A, \lambda_E, \lambda_F^{1:M}\right)\right] P\left(f_D\right) P\left(f_A\right). \end{aligned} \tag{S.51}$$

As $f_D$ and $f_A$ both are binary variables, we place Bernoulli priors on them (Eqs. (S.39) and (S.40)). Therefore, we get two Bernoulli posteriors dependent upon the observation likelihood specified in Eqs. (S.47) to (S.49)

$$\begin{aligned} &f_D\middle|d_{1:K}, \delta_{1:K}, c_{1:K}, \mathbf{x}_{1:K}, f_A, \lambda_E, \lambda_F^{1:M} \\ &\quad \sim \mathbf{Bernoulli}\left(\left[1 + \prod_{k=1}^{K} \frac{P\left(d_k, \delta_k\middle|c_k, \mathbf{x}_k, f_D = 0, f_A, \lambda_E, \lambda_F^{1:M}\right)}{P\left(d_k, \delta_k\middle|c_k, \mathbf{x}_k, f_D = 1, f_A, \lambda_E, \lambda_F^{1:M}\right)}\right]^{-1}\right), \end{aligned} \tag{S.52}$$

$$\begin{aligned} &f_A\middle|d_{1:K}, \delta_{1:K}, c_{1:K}, \mathbf{x}_{1:K}, f_D, \lambda_E, \lambda_F^{1:M} \\ &\quad \sim \mathbf{Bernoulli}\left(\left[1 + \prod_{k=1}^{K} \frac{P\left(d_k, \delta_k\middle|c_k, \mathbf{x}_k, f_D, f_A = 0, \lambda_E, \lambda_F^{1:M}\right)}{P\left(d_k, \delta_k\middle|c_k, \mathbf{x}_k, f_D, f_A = 1, \lambda_E, \lambda_F^{1:M}\right)}\right]^{-1}\right). \end{aligned} \tag{S.53}$$

#### C.3 Learning spatial trajectory

Similarly, the terms in Eq. (S.50) with dependence upon $\mathbf{x}_{1:K}$ form the conditional posterior of spatial trajectories:

$$
\begin{aligned}
&P\left(\mathbf{x}_{1:K}\middle|d_{1:K}, \delta_{1:K}, c_{1:K}, f_D, f_A, \mathbf{\Pi}, D_{1:M}, \lambda_E, \lambda_F^{1:M}\right)\\
&\propto \left[\prod_{k=1}^{K} P\left(d_k\middle|c_k, \mathbf{x}_k, f_D, f_A, \lambda_E, \lambda_F^{1:M}\right)\right]\left[\prod_{k=1}^{K-1} P\left(\mathbf{x}_{k+1}|c_k, \mathbf{x}_k, D_{1:M}\right)\right] P\left(\mathbf{x}_1\right) \qquad \text{(S.54)}\\
&\propto \left(\prod_{k=1}^{K} P_{d_k=\varnothing}^{\delta_{\varnothing,d_k}} P_{d_k=D}^{\delta_{1,d_k}} P_{d_k=A}^{\delta_{2,d_k}}\right)\left[\prod_{k=1}^{K-1} \textbf{Normal}\left(\mathbf{x}_{k+1}; \mathbf{x}_k, 2D_{c_k}T\right)\right]\\
&\quad\times \textbf{Normal}\left(x_1; x_0, \sigma_x^2\right) \textbf{Normal}\left(y_1; y_0, \sigma_y^2\right) \textbf{Normal}\left(z_1; z_0, \sigma_z^2\right) \qquad \text{(S.55)}\\
&\propto \left(\prod_{k=1}^{K} P_{d_k=\varnothing} p_{d_k=D}^{\delta_{1,d_k}} p_{d_k=A}^{\delta_{2,d_k}}\right)\left[\prod_{k=1}^{K-1} \textbf{Normal}\left(\mathbf{x}_{k+1}; \mathbf{x}_k, 2D_{c_k}T\right)\right]\\
&\quad\times \textbf{Normal}\left(x_1; x_0, \sigma_x^2\right) \textbf{Normal}\left(y_1; y_0, \sigma_y^2\right) \textbf{Normal}\left(z_1; z_0, \sigma_z^2\right). \qquad \text{(S.56)}
\end{aligned}
$$

Here, we've chosen a Normal distribution as the prior on the initial location, and $p_{d_k=D} = P_{d_k=D}/P_{d_k=\varnothing}$.

##### C.3.1 Hamiltonian Monte Carlo

A traditional means of trajectory sampling would be by filtering, however, on account of the nonlinearity of the likelihood over position, we would require a nonlinear filter. As an alternative to nonlinear filter which would require number of approximations here we use Hamilton Monte Carlo (HMC) [3]. Briefly, HMC treats the logarithm of a probability density function (which is our conditional posterior) as an energy landscape and samples are proposed according to Hamiltonian mechanics. Thus, 2e start from calculating the negative log posterior of the spatial trajectory,

$$
\begin{aligned}
&-\ln P\left(\mathbf{x}_{1:K}\middle|d_{1:K}, \delta_{1:K}, c_{1:K}, f_D, f_A, \mathbf{\Pi}, D_{1:M}, \lambda_E, \lambda_F^{1:M}\right)\\
&= \text{Const.} - \sum_{k=1}^{K}\left(\ln P_{d_k=\varnothing} + \delta_{1,d_k}\ln p_{d_k=D} + \delta_{2,d_k}\ln p_{d_k=A}\right) + \sum_{k=1}^{K-1}\frac{\left(\mathbf{x}_{k+1}-\mathbf{x}_k\right)^2}{4D_{c_k}T}\\
&\quad+\frac{(x_1-x_0)^2}{2\sigma_x^2} + \frac{(y_1-y_0)^2}{2\sigma_y^2} + \frac{(z_1-z_0)^2}{2\sigma_z^2}. \qquad \text{(S.57)}
\end{aligned}
$$

Now we can define

$$V(\mathbf{x}_{1:K}) = -\sum_{k=1}^{K}\left(\ln P_{d_k=\varnothing} + \delta_{1,d_k}\ln p_{d_k=D} + \delta_{2,d_k}\ln p_{d_k=A}\right), \qquad \text{(S.58)}$$

$$L(\mathbf{x}_{1:K}) = \frac{(x_1-x_0)^2}{2\sigma_x^2} + \frac{(y_1-y_0)^2}{2\sigma_y^2} + \frac{(z_1-z_0)^2}{2\sigma_z^2} + \sum_{k=1}^{K-1}\frac{\left(\mathbf{x}_{k+1}-\mathbf{x}_k\right)^2}{4D_{c_k}T}. \qquad \text{(S.59)}$$

Here, for illustrative purposes, we show how we sample the $x$-component. In order to implement the method of HMC, the full Hamiltonian of the system can be written as

$$\mathcal{H}(x_{1:K}, p_{1:K}) = T(p_{1:K}) + V(x_{1:K}) + L(x_{1:K}), \tag{S.60}$$

where $p_{1:K}$ are generalized momenta and the "kinetic energy" $T(p_{1:K})$ is defined as

$$T(p_{1:K}) = \frac{1}{2} p_{1:K}^T \mathbf{M}^{-1} p_{1:K} \tag{S.61}$$

with

$$\mathbf{M} = \begin{bmatrix} m_1 & 0 & \cdot & 0 \\ 0 & m_2 & \cdot & 0 \\ \vdots & \vdots & \ddots & \vdots \\ 0 & 0 & \cdot & m_K \end{bmatrix}, \quad p_{1:K} = \begin{bmatrix} p_1 \\ p_2 \\ \vdots \\ p_K \end{bmatrix}. \tag{S.62}$$

From Hamiltonian mechanics we know that the time evolution of the system is dictated by

$$\begin{cases} \dfrac{\mathrm{d}x_k}{\mathrm{d}t} = \dfrac{\partial \mathcal{H}}{\partial p_k}, \\ \dfrac{\mathrm{d}p_k}{\mathrm{d}t} = -\dfrac{\partial \mathcal{H}}{\partial x_k}. \end{cases} \tag{S.63}$$

Unfortunately, Eq. (S.63) cannot be solved analytically here and some numerical recipe must be applied.

In order to numerically evolve $\mathcal{H}$ over "time" with high precision, we invoke Strang splitting [4], whose error is $\mathcal{O}(h^2)$ with $h$ being the step size in "time". This is done by first split the original Hamiltonian into two parts

$$\mathcal{H}(x_{1:K}, p_{1:K}) = [V(p_{1:K})] + [T(p_{1:K}) + L(p_{1:K})] \tag{S.64}$$

$$= \mathcal{H}^1(x_{1:K}) + \mathcal{H}^2(x_{1:K}, p_{1:K}). \tag{S.65}$$

Equipped with the two equations above, we can now provide a full algorithm below:

1. Advance a half-step using $\mathcal{H}^1(x_{1:K})$,
$$x_{1:K}^i \to x_{1:K}^{int,1}, \; p_{1:K}^i \to p_{1:K}^{int,1};$$
2. Advance a whole-step using $\mathcal{H}^2(x_{1:K}, p_{1:K})$,
$$x_{1:K}^{int,1} \to x_{1:K}^{int,2}, \; p_{1:K}^{int,1} \to p_{1:K}^{int,2};$$
3. Advance a half-step using $\mathcal{H}^1(x_{1:K})$,
$$x_{1:K}^{int,2} \to x_{1:K}^{f}, \; p_{1:K}^{int,2} \to p_{1:K}^{f}.$$

Normally, we should require the Hamiltonian to be conserved to infinite precision. However, this cannot be achieved for any numerical calculation. Thus, we calculate an acceptance ratio based on the difference between Hamiltonians before and after the numerical integration. And this ratio is used to determine whether the proposed spatial trajectory sample should be accepted.

##### C.3.2 Renormalization of $x$ and $p$

Before proceeding further, for computational simplicity, we can renormalize the positions with respect to the confocal region's dimensions,

$$X_{1:K} = \frac{1}{w_x} x_{1:K}. \tag{S.66}$$

Although Eq. (S.58) stays the same, Eq. (S.59) does require some modification. In addition, when $x$ is being updated, neither $y$ nor $z$ should be changed.

Therefore, the part of the "energy" not depending on $x$ is always conserved, and hence, does not need to be considered,

$$L_x(X_{1:K}) = \frac{(X_1 - X_0)^2}{2\sigma_X^2} + \sum_{k=1}^{K-1} \left[ \frac{w_x^2}{4D_{c_k}T} (X_{k+1} - X_k)^2 \right]. \tag{S.67}$$

Also note that generalized momenta $P_{1:K}$ is used in correspondence to $X_{1:K}$.

##### C.3.3 Advance with $\mathcal{H}^1$

After the aforementioned renormalization, we continue describing the full algorithm of Strang splitting. When we advance the system with $\mathcal{H}^1(x_{1:K})$, Eq. (S.63) becomes

$$\begin{cases} \dfrac{\mathrm{d}X_k}{\mathrm{d}t} = \dfrac{\partial \mathcal{H}^1}{\partial P_k} = 0, \\ \dfrac{\mathrm{d}P_k}{\mathrm{d}t} = -\dfrac{\partial \mathcal{H}^1}{\partial X_k} = -\dfrac{\partial V}{\partial X_k}. \end{cases} \tag{S.68}$$

Consequently,

$$\begin{cases} X_k^{int,1} = X_k^i, \\ P_k^{int,1} = P_k^i - \dfrac{h}{2} \dfrac{\partial V}{\partial X_k} \Big|_{X_k^i}, \end{cases} \tag{S.69}$$

and

$$\begin{cases} X_k^f = X_k^{int,2}, \\ P_k^f = P_k^{int,2} - \dfrac{h}{2} \dfrac{\partial V}{\partial X_k} \Big|_{X_k^{int,2}}. \end{cases} \tag{S.70}$$

The derivative of $V$ is given by

$$\frac{\partial V}{\partial X_k} = - \frac{\partial}{\partial X_k} \left( \ln P_{d_k=\varnothing} + \delta_{1,d_k} \ln P_{d_k=D} + \delta_{2,d_k} \ln P_{d_k=A} \right) \tag{S.71}$$

$$= - \left( \frac{1}{P_{d_k=\varnothing}} \frac{\partial P_{d_k=\varnothing}}{\partial X_k} + \frac{\delta_{1,d_k}}{P_{d_k=D}} \frac{\partial p_{d_k=D}}{\partial X_k} + \frac{\delta_{2,d_k}}{p_{d_k=A}} \frac{\partial P_{d_k=A}}{\partial X_k} \right). \tag{S.72}$$

$$
\begin{aligned}
\frac{\partial P_{d_k=\varnothing}}{\partial X_k} =& \frac{\partial}{\partial X_k}\left(e^{\mu_{E,k}^D \tau} + e^{\mu_{E,k}^A \tau} + e^{\lambda_B^D T} + e^{\lambda_B^A T} - 3\right)^{-1} && \text{(S.73)}\\
=& -\tau P_{d_k=\varnothing}^2 \left(\mu_{E,k}^D e^{\mu_{E,k}^D \tau} + \mu_{E,k}^A e^{\mu_{E,k}^A \tau}\right) \frac{\partial \mathrm{PSF}_k}{\partial X_k}, && \text{(S.74)}\\
\frac{\partial p_{d_k=D}}{\partial X_k} =& \tau\left(\mu_{E,k}^D e^{\mu_{E,k}^D \tau} \frac{\mu_R^D P_{DD} + \mu_F^{c_k} P_{AD}}{\mu_R^D + \mu_F^{c_k}} + \mu_{E,k}^A e^{\mu_{E,k}^A \tau} P_{AD}\right) \frac{\partial \mathrm{PSF}_k}{\partial X_k}, && \text{(S.75)}\\
\frac{\partial p_{d_k=A}}{\partial X_k} =& \tau\left(\mu_{E,k}^D e^{\mu_{E,k}^D \tau} \frac{\mu_R^D P_{DA} + \mu_F^{c_k} P_{AA}}{\mu_R^D + \mu_F^{c_k}} + \mu_{E,k}^A e^{\mu_{E,k}^A \tau} P_{AA}\right) \frac{\partial \mathrm{PSF}_k}{\partial X_k}. && \text{(S.76)}
\end{aligned}
$$

$\frac{\partial \mathrm{PSF}_k}{\partial X_k}$ needs to be calculated based on the actual PSF used which, for a three-dimensional Gaussian, reads

$$
\frac{\partial \mathrm{PSF}_k}{\partial X_k} = \frac{\partial}{\partial X_k} \exp\left[-2\left(X_k^2 + Y_k^2 + Z_k^2\right)\right] = -4X_k \mathrm{PSF}_k. \tag{S.77}
$$

Note that for other types of PSF, we would simply insert the corresponding expression into Eq. (S.77).

##### C.3.4 Advance with $\mathcal{H}^2$

The next step in Strang splitting is to advance the system using $\mathcal{H}^2(X_{1:K}, P_{1:K})$,

$$
\begin{cases}
\frac{\mathrm{d}X_{1:K}}{\mathrm{d}t} = \frac{\partial \mathcal{H}^2}{\partial P_{1:K}} = \frac{\partial T}{\partial P_{1:K}} = \mathbf{M}^{-1} P_{1:K},\\
\frac{\mathrm{d}P_{1:K}}{\mathrm{d}t} = -\frac{\partial \mathcal{H}^2}{\partial X_{1:K}} = -\frac{\partial L}{\partial X_{1:K}}.
\end{cases} \tag{S.78}
$$

Then from Eq. (S.67),

$$\frac{\partial L}{\partial X_{1:K}} \tag{S.79}$$

$$=w_x^2 \begin{bmatrix} \frac{X_1}{\sigma_X^2 w_x^2} + \frac{X_1}{2D_{c_1}T} - \frac{X_2}{2D_{c_1}T} \\ -\frac{X_1}{2D_{c_1}T} + \frac{X_2}{2D_{c_1}T} + \frac{X_2}{2D_{c_2}T} - \frac{X_3}{2D_{c_2}T} \\ \vdots \\ -\frac{X_{K-2}}{2D_{c_{K-2}}T} + \frac{X_{K-1}}{2D_{c_{K-2}}T} + \frac{X_{K-1}}{2D_{c_{K-1}}T} - \frac{X_K}{2D_{c_{K-1}}T} \\ -\frac{X_{K-1}}{2D_{c_{K-1}}T} + \frac{X_K}{2D_{c_{K-1}}T} \end{bmatrix} - \begin{bmatrix} \frac{X_0}{\sigma_X^2} \\ 0 \\ \vdots \\ 0 \\ 0 \end{bmatrix} \tag{S.80}$$

$$=\frac{w_x^2}{2T} \begin{bmatrix} \frac{2T}{\sigma_X^2 w_x^2} + D_{c_1}^{-1} & -D_{c_1}^{-1} & & & 0 \\ -D_{c_1}^{-1} & D_{c_1}^{-1} + D_{c_2}^{-1} & -D_{c_2}^{-1} & & \\ & -D_{c_2}^{-1} & D_{c_2}^{-1} + D_{c_3}^{-1} & \ddots & \\ & & \ddots & \ddots & -D_{c_{K-1}}^{-1} \\ 0 & & & -D_{c_{K-1}}^{-1} & D_{c_{K-1}}^{-1} \end{bmatrix} \begin{bmatrix} X_1 \\ X_2 \\ \vdots \\ X_{K-1} \\ X_K \end{bmatrix} - \begin{bmatrix} \frac{X_0}{\sigma_X^2} \\ 0 \\ \vdots \\ 0 \\ 0 \end{bmatrix} \tag{S.81}$$

$$=\frac{w_x^2}{2T}\mathbf{A}X_{1:K} - \nu_X, \tag{S.82}$$

where $\mathbf{A}$ and $\nu$ are direct substitutions. It follows that

$$\begin{cases} X_{1:K}^{int,2} - X_{1:K}^{int,1} = h\mathbf{M}^{-1}\frac{P_{1:K}^{int,1} + P_{1:K}^{int,2}}{2}, \\ P_{1:K}^{int,2} - P_{1:K}^{int,1} = -\frac{hw_x^2}{2T}\mathbf{A}\frac{X_{1:K}^{int,1} + X_{1:K}^{int,2}}{2} + h\nu_X. \end{cases} \tag{S.83}$$

The solution to this linear system is given by

$$\left(\frac{2}{h}\mathbf{M} + \frac{hw_x^2}{4T}\mathbf{A}\right) X_{1:K}^{int,2} = \left(\frac{2}{h}\mathbf{M} - \frac{hw_x^2}{4T}\mathbf{A}\right) X_{1:K}^{int,1} + 2P_{1:K}^{int,1} + h\nu_X, \tag{S.84}$$

$$P_{1:K}^{int,2} = \frac{2\mathbf{M}}{h}\left(X_{1:K}^{int,2} - X_{1:K}^{int,1}\right) - P_{1:K}^{int,1}. \tag{S.85}$$

For the convenience of further discussion, let

$$f_{1:K} = \left(\frac{2}{h}\mathbf{M} - \frac{hw_x^2}{4T}\mathbf{A}\right) X_{1:K}^{int,1} + 2P_{1:K}^{int,1} + h\nu_X, \tag{S.86}$$

then Eq. (S.84) becomes

$$\left(\frac{2}{h}\mathbf{M} + \frac{hw_x^2}{4T}\mathbf{A}\right) X_{1:K} = f_{1:K}. \tag{S.87}$$

Recall that $\mathbf{M}$ is diagonal and $\mathbf{A}$ is tri-diagonal, as such, $\frac{2}{h}\mathbf{M} + \frac{hw_x^2}{4T}\mathbf{A}$ must also be a tri-diagonal matrix.

Clearly, in order to obtain $X_{1:K}$, we must invert $\frac{2}{h}\mathbf{M} + \frac{hw_x^2}{4T}\mathbf{A}$. As $K$ can be a large number, calculating the inverse matrix using the naive Gauss-Jordan elimination can be computationally heavy $\mathcal{O}(K^3)$. However, this issue can be greatly resolved. Recall that $\frac{2}{h}\mathbf{M} + \frac{hw_x^2}{4T}\mathbf{A}$ is a tri-diagonal matrix, this feature allows us to use the Thomas algorithm, whose cost is $\mathcal{O}(K)$.

To demonstrate this method, we can perform the substitution that

$$\frac{2}{h}\mathbf{M} + \frac{hw_x^2}{4T}\mathbf{A} = \begin{bmatrix} \phi_1 & \xi_1 & & & 0 \\ \xi_1 & \phi_2 & \xi_2 & & \\ & \xi_2 & \phi_3 & \ddots & \\ & & \ddots & \ddots & \xi_{K-1} \\ 0 & & & \xi_{K-1} & \phi_K \end{bmatrix} \tag{S.88}$$

with

$$\xi_k = -\frac{hw_x^2}{4T} D_{c_k}^{-1}, \tag{S.89}$$

$$\phi_k = \frac{2m_n}{h} + \frac{h}{2} \begin{cases} \sigma_X^{-2} + \dfrac{w_x^2}{2T} D_{c_1}^{-1} & \text{for } k = 1; \\ \dfrac{w_x^2}{2T}\left(D_{c_{k-1}}^{-1} + D_{c_k}^{-1}\right) & \text{for } k = 2, 3, \ldots, K-1; \\ \dfrac{w_x^2}{2T} D_{c_{K-1}}^{-1} & \text{for } k = K. \end{cases} \tag{S.90}$$

As an implementation Thomas algorithm, we must first march forward as follows

$$\xi'_k = \begin{cases} \dfrac{\xi_1}{\phi_1} & \text{for } k = 1; \\ \dfrac{\xi_k}{\phi_k - \xi_{k-1}\xi'_{k-1}} & \text{for } k = 2, 3, \ldots, K-1, \end{cases} \tag{S.91}$$

$$f'_k = \begin{cases} \dfrac{f_1}{\phi_1} & \text{for } k = 1; \\ \dfrac{f_k - \xi_{k-1} f'_{k-1}}{\phi_k - \xi_{k-1}\xi'_{k-1}} & \text{for } k = 2, 3, \ldots, K, \end{cases} \tag{S.92}$$

and by marching backward we have,

$$X_k^{int,2} = \begin{cases} f'_k - \xi'_k X_{k+1}^{int,2} & \text{for } k = 1, 2, \ldots, K-1; \\ f'_K & \text{for } k = K. \end{cases} \tag{S.93}$$

##### C.3.5 Summary of Spatial Trajectory HMC

For the sake of easier reference and to refresh readers' memory, we summarize some key equations in this section:

$$\begin{cases} X_k^{int,1} = X_k^i, \\ P_k^{int,1} = P_k^i - \dfrac{h}{2}\dfrac{\partial V}{\partial X_k}\Big|_{X_k^i}, \end{cases} \tag{S.94}$$

$$\begin{cases} X_k^{int,2} = \begin{cases} f'_k - \xi'_k X_{k+1}^{int,2} & \text{for } k = 1, 2, \dots, K-1; \\ f'_K & \text{for } k = K. \end{cases} \\ P_{1:K}^{int,2} = \dfrac{2\mathbf{M}}{h}\left(X_{1:K}^{int,2} - X_{1:K}^{int,1}\right) - P_{1:K}^{int,1} \end{cases} \tag{S.95}$$

$$\begin{cases} X_k^f = X_k^{int,2}, \\ P_k^f = P_k^{int,2} - \dfrac{h}{2}\dfrac{\partial V}{\partial X_k}\Big|_{X_k^{int,2}}. \end{cases} \tag{S.96}$$

#### C.4 Learning states

Once the spatial trajectorie is updated, we move on to the third step of our Gibbs scheme described in Section C.1, sampling the state trajectory. The target distribution is $P\left(c_{1:K}\middle|d_{1:K}, \delta_{1:K}, \mathbf{x}_{1:K}, f_D, f_A,\right.$ and it can be expanded as

$$\begin{aligned} &P\left(c_{1:K}\middle|d_{1:K}, \delta_{1:K}, \mathbf{x}_{1:K}, f_D, f_A, \mathbf{\Pi}, D_{1:M}, \lambda_E, \lambda_F^{1:M}\right) \\ &= P\left(c_1\middle|c_2, d_{1:K}, \delta_{1:K}, \mathbf{x}_{1:K}, f_D, f_A, \mathbf{\Pi}, D_{1:M}, \lambda_E, \lambda_F^{1:M}\right) \\ &\times P\left(c_2\middle|c_3, d_{1:K}, \delta_{1:K}, \mathbf{x}_{1:K}, f_D, f_A, \mathbf{\Pi}, D_{1:M}, \lambda_E, \lambda_F^{1:M}\right) \\ &\qquad\vdots \\ &\times P\left(c_{K-1}\middle|c_K, d_{1:K}, \delta_{1:K}, \mathbf{x}_{1:K}, f_D, f_A, \mathbf{\Pi}, D_{1:M}, \lambda_E, \lambda_F^{1:M}\right) \\ &\times P\left(c_K\middle|d_{1:K}, \delta_{1:K}, \mathbf{x}_{1:K}, f_D, f_A, \mathbf{\Pi}, D_{1:M}, \lambda_E, \lambda_F^{1:M}\right). \end{aligned} \tag{S.97}$$

Here, we define the forward filter

$$\mathcal{A}_k\left(c_k\right) \equiv P\left(c_k\middle|d_{1:K}, \delta_{1:K}, \mathbf{x}_{1:K}, f_D, f_A, \mathbf{\Pi}, D_{1:M}, \lambda_E, \lambda_F^{1:M}\right). \tag{S.98}$$

Now the last term Eq. (S.97), becomes $\mathcal{A}_K\left(c_K\right)$, and the rest can also be expressed in terms of filters. For $k = 1, 2 \dots, K-1$,

$$\begin{aligned} &P\left(c_k\middle|c_{k+1}, d_{1:K}, \delta_{1:K}, \mathbf{x}_{1:K}, f_D, f_A, \mathbf{\Pi}, D_{1:M}, \lambda_E, \lambda_F^{1:M}\right) \\ &\propto P\left(c_{k+1}\middle|c_k, d_{1:K}, \delta_{1:K}, \mathbf{x}_{1:K}, f_D, f_A, \mathbf{\Pi}, D_{1:M}, \lambda_E, \lambda_F^{1:M}\right) \\ &\times P\left(c_k\middle|d_{1:K}, \delta_{1:K}, \mathbf{x}_{1:K}, f_D, f_A, \mathbf{\Pi}, D_{1:M}, \lambda_E, \lambda_F^{1:M}\right) && \text{(S.99)} \\ &\propto P\left(c_{k+1}|c_k, \mathbf{\Pi}\right) P\left(c_k\middle|d_{1:K}, \delta_{1:K}, \mathbf{x}_{1:K}, f_D, f_A, \mathbf{\Pi}, D_{1:M}, \lambda_E, \lambda_F^{1:M}\right) && \text{(S.100)} \\ &\propto \pi_{c_k, c_{k+1}}\mathcal{A}_k\left(c_k\right). && \text{(S.101)} \end{aligned}$$

Therefore, the state trajectory posterior probability distribution is turned to a product of transition probabilities and filters,

$$P\left(c_{1:K}\middle|d_{1:K},\delta_{1:K},\mathbf{x}_{1:K},f_D,f_A,\mathbf{\Pi},D_{1:M},\lambda_E,\lambda_F^{1:M}\right)\propto\left[\prod_{k=1}^{K-1}\pi_{c_k,c_{k+1}}\mathcal{A}_k\left(c_k\right)\right]\mathcal{A}_K\left(c_K\right). \tag{S.102}$$

All that is left to do is constructing all the filters recursively. For any $k=2,...,K-1$,

$$\begin{aligned}\mathcal{A}_k\left(c_k\right)&=\sum_{c_{k-1}}P\left(c_k,c_{k-1}\middle|d_{1:K},\delta_{1:K},\mathbf{x}_{1:K},f_D,f_A,\mathbf{\Pi},D_{1:M},\lambda_E,\lambda_F^{1:M}\right) &\text{(S.103)}\\&=\sum_{c_{k-1}}P\left(c_k\middle|c_{k-1},d_{1:K},\delta_{1:K},\mathbf{x}_{1:K},f_D,f_A,\mathbf{\Pi},D_{1:M},\lambda_E,\lambda_F^{1:M}\right)\\&\quad\times P\left(c_{k-1}\middle|d_{1:K},\delta_{1:K},\mathbf{x}_{1:K},f_D,f_A,\mathbf{\Pi},D_{1:M},\lambda_E,\lambda_F^{1:M}\right) &\text{(S.104)}\\&=\sum_{c_{k-1}}P\left(c_k\middle|c_{k-1},d_{1:K},\delta_{1:K},\mathbf{x}_{1:K},f_D,f_A,\mathbf{\Pi},D_{1:M},\lambda_E,\lambda_F^{1:M}\right)\mathcal{A}_{k-1}\left(c_{k-1}\right). &\text{(S.105)}\end{aligned}$$

From the direct comparison with Eq. (S.50), we know

$$\begin{aligned}&P\left(c_k\middle|c_{k-1},d_{1:K},\delta_{1:K},\mathbf{x}_{1:K},f_D,f_A,\mathbf{\Pi},D_{1:M},\lambda_E,\lambda_F^{1:M}\right)\\&\propto P\left(d_k,\delta_k\middle|c_k,\mathbf{x}_k,f_D,f_A,\lambda_E,\lambda_F^{1:M}\right)P\left(\mathbf{x}_{k+1}|c_k,\mathbf{x}_k,D_{1:M}\right)P\left(c_k|c_{k-1},\mathbf{\Pi}\right), \end{aligned}\tag{S.106}$$

so

$$\begin{aligned}&\mathcal{A}_k\left(c_k\right)\\&\propto P\left(d_k,\delta_k\middle|c_k,\mathbf{x}_k,f_D,f_A,\lambda_E,\lambda_F^{1:M}\right)P\left(\mathbf{x}_{k+1}|c_k,\mathbf{x}_k,D_{1:M}\right)\sum_{c_{k-1}}\pi_{c_{k-1},c_k}\mathcal{A}_{k-1}\left(c_{k-1}\right).\end{aligned}\tag{S.107}$$

By including $k=1$ and $k=K$ we have

$$\begin{aligned}\mathcal{A}_1\left(c_1\right)&\propto P\left(d_1,\delta_1\middle|c_1,\mathbf{x}_1,f_D,f_A,\lambda_E,\lambda_F^{1:M}\right)P\left(\mathbf{x}_2|\mathbf{x}_1,c_1,D_{1:M}\right)P\left(c_1\right), &\text{(S.108)}\\\mathcal{A}_k\left(c_k\right)&\propto P\left(d_k,\delta_k\middle|c_k,\mathbf{x}_k,f_D,f_A,\lambda_E,\lambda_F^{1:M}\right)P\left(\mathbf{x}_{k+1}|c_k,\mathbf{x}_k,D_{1:M}\right)\\&\qquad\times\sum_{c_{k-1}}\pi_{c_{k-1},c_k}\mathcal{A}_{k-1}\left(c_{k-1}\right), &\text{(S.109)}\\\mathcal{A}_K\left(c_K\right)&\propto P\left(d_K,\delta_K\middle|c_K,\mathbf{x}_K,f_D,f_A,\lambda_E,\lambda_F^{1:M}\right)\sum_{c_{K-1}}\pi_{c_{K-1},c_K}\mathcal{A}_{K-1}\left(c_{K-1}\right). &\text{(S.110)}\end{aligned}$$

Note that the proportionality can be obtained by requiring $\sum_{c_k}\mathcal{A}_k\left(c_k\right)=1$. Finally, the entire trajectory can be sampled from

$$c_k\middle|d_K,\delta_K,c_{k+1},\mathbf{x}_K,f_D,f_A,\lambda_E,\lambda_F^{1:M}\sim\mathbf{Categorical}_{1:M}\left(P_{c_k=1},P_{c_k=2},\dots,P_{c_k=M}\right) \tag{S.111}$$

$$P_{c_k=m}=\begin{cases}\mathcal{A}_k(m), & k=K;\\ \pi_{m,c_{k+1}}\mathcal{A}_k(m)/\sum_{m=1}^{M}\pi_{m,c_{k+1}}\mathcal{A}_k(m), & \text{otherwise.}\end{cases} \tag{S.112}$$

#### C.5 Learning transition probabilities

The target distribution is $P(\mathbf{\Pi}|c_{1:K})$, according to the Bayesian theorem,

$$P(\mathbf{\Pi}|c_{1:K}) \propto P(c_{2:K}|c_1, \mathbf{\Pi}) P(\mathbf{\Pi}) = \left(\prod_{k=1}^{K-1} \pi_{c_k,c_{k+1}}\right) P(\mathbf{\Pi}). \tag{S.113}$$

If we denote the number of transitions from state $\sigma_1$ to state $\sigma_2$ as $n_{1,2}$, the equation above then becomes

$$P(\mathbf{\Pi}|c_{1:K}) \propto \pi_{\sigma_1,\sigma_1}^{n_{1,1}} \cdots \pi_{\sigma_M,\sigma_M}^{n_{M,M}} P(\mathbf{\Pi}) \tag{S.114}$$

$$\propto \prod_{m=1}^{M} \pi_{\sigma_m,\sigma_1}^{n_{m,1}} \cdots \pi_{\sigma_m,\sigma_M}^{n_{m,M}} P(\boldsymbol{\pi}_m|\bar{\alpha}_{\pi,m}). \tag{S.115}$$

In fact, for each $m$, $P(\boldsymbol{\pi}_m|c_{1:K}) \propto \pi_{\sigma_m,\sigma_1}^{n_{m,1}} \cdots \pi_{\sigma_m,\sigma_M}^{n_{m,M}} P(\boldsymbol{\pi}_m|\bar{\alpha}_{\pi,m})$. More accurately,

$$\boldsymbol{\pi}_m|c_{1:K} \sim \mathbf{Dirichlet}(\bar{n}_m + \bar{\alpha}_{\pi,m}) \tag{S.116}$$

where $\bar{n}_m = (n_{m,1}, \ldots, n_{m,M})$.

#### C.6 Learning diffusion coefficients

The posterior on the diffusion coefficient of the $m$th state is $P(D_m|c_{1:K}, \mathbf{x}_{1:K}, \alpha_{D_m}, \beta_{D_m})$. It follows that

$$P(D_m|c_{1:K}, \mathbf{x}_{1:K}, \alpha_{D_m}, \beta_{D_m})$$

$$= P(D_m|\{\mathbf{x}_k, \mathbf{x}_{k+1}|c_k = m\}, \alpha_{D_m}, \beta_{D_m}) \tag{S.117}$$

$$\propto P(D_m, \{\mathbf{x}_{k+1}|c_k = m\}|\{\mathbf{x}_k|c_k = m\}, \alpha_{D_m}, \beta_{D_m}) \tag{S.118}$$

$$\propto P(\{\mathbf{x}_{k+1}|c_k = m\}|D_m, \{\mathbf{x}_k|c_k = m\}) P(D_m|, \alpha_{D_m}, \beta_{D_m}). \tag{S.119}$$

Here, $P(D_m|\cdot)$ is the prior on the $m$th diffusion coefficient. Furthermore, the likelihood part of Eq. (S.119) can be written as

$$P(\{\mathbf{x}_{k+1}|c_k = m\}|D_m, \{\mathbf{x}_k|c_k = m\}) = \prod_{\substack{c_k=m \\ k<K}} P(\mathbf{x}_{k+1}|D_m, \mathbf{x}_k) \tag{S.120}$$

$$= \prod_{\substack{c_k=m \\ k<K}} \mathbf{Normal}(\mathbf{x}_{k+1}; \mathbf{x}_k, 2D_mT). \tag{S.121}$$

Letting $\Delta\mathbf{x}_k \equiv \mathbf{x}_{k+1} - \mathbf{x}_k$ simplifies $\mathbf{Normal}(\mathbf{x}_{k+1}; \mathbf{x}_k, 2D_mT)$ to $\mathbf{Normal}(\Delta\mathbf{x}_k; \mathbf{0}, 2D_mT)$. By assuming $k_m$ is the index of the well-ordered set $\{\Delta\mathbf{x}_k|c_k = m\}$ and $K_m$ is the order of this set we can further write

$$\prod_{\substack{c_k=m \\ k<K}} \mathbf{Normal}(\mathbf{x}_k, 2D_mT) = \left(\frac{1}{4\pi D_mT}\right)^{3K_m/2} \exp\left(-\frac{\sum_{k_m=1}^{K_m} \Delta\mathbf{x}_{k_m}^2}{4D_mT}\right). \tag{S.122}$$

Consequently,

$$\begin{aligned} &P\left(D_m|c_{1:K},\mathbf{x}_{1:K}\right) \\ &\propto P\left(\{\mathbf{x}_{k+1}|c_k=m\}|D_m,\{\mathbf{x}_k|c_k=m\}\right)P\left(D_m|\alpha_{D_m},\beta_{D_m}\right) &\text{(S.123)} \\ &\propto \left(\frac{1}{D_m}\right)^{\alpha_{D_m}+1+3K_m/2}\exp\left(-\frac{\sum_{k_m=1}^{K_m}\Delta\mathbf{x}_{k_m}^2}{4D_mT}-\frac{\beta_{D_m}}{D_m}\right) &\text{(S.124)} \\ &\propto \textbf{Inv-Gamma}\left(D_m;\alpha_{D_m}+\frac{3K_m}{2},\beta_{D_m}+\sum_{k_m=1}^{K_m}\frac{\Delta\mathbf{x}_{k_m}^2}{4T}\right). &\text{(S.125)} \end{aligned}$$

#### C.7 Learning FRET rates

The posterior on the FRET rate of the $m$th state is

$$\begin{aligned} &P\left(\lambda_F^m|d_{1:K},\delta_{1:K},c_{1:K},\mathbf{x}_{1:K},f_D,f_A,\lambda_E\right) \\ &\propto P\left(\delta_{1:K},d_{1:K}|c_{1:K},\mathbf{x}_{1:K},f_D,f_A,\lambda_E,\lambda_F^m\right)P\left(\lambda_F^m\right). &\text{(S.126)} \end{aligned}$$

Since the process of FRET only comes into play when a signal photon is emitted ($o_k=1$) and $\lambda_F^m$ is involved only if a molecule is in its unbound state ($c_k=1$), the form of posterior can be simplified to

$$P\left(\lambda_F^m|d_{1:K},\delta_{1:K},c_{1:K},\mathbf{x}_{1:K},f_D,f_A,\lambda_E\right)\propto P\left(\delta_{1:\mathcal{K}_1},e_{1:\mathcal{K}_1}|\lambda_F^m\right)P\left(\lambda_F^m\right), \tag{S.127}$$

where $\kappa_1$ is the index of signal photons emitted in the unbound state and $\mathcal{K}_1$ is the total number of those.

The first piece is the likelihood regarding the arrival time and detection channel,

$$P\left(\delta_{1:\mathcal{K}_1},e_{1:\mathcal{K}_1}|\lambda_F^m\right)=\prod_{\kappa_1=1}^{\mathcal{K}_1}P\left(\delta_{\kappa_1},e_{\kappa_1}|\lambda_F^m\right). \tag{S.128}$$

As for the prior part we choose the Gamma distribution

$$P\left(\lambda_F^m\right)=\textbf{Gamma}\left(\alpha_F^U,\frac{\beta_F^U}{\alpha_F^U}\right), \tag{S.129}$$

so

$$P\left(\lambda_F^m|d_{1:K},\delta_{1:K},c_{1:K},\mathbf{x}_{1:K},f_D,f_A,\lambda_E\right)\propto\left[\prod_{\kappa_1=1}^{\mathcal{K}_1}P\left(\delta_{\kappa_1},e_{\kappa_1}|\lambda_F^m\right)\right]\textbf{Gamma}\left(\lambda_F^m;\alpha_F^U,\frac{\beta_F^U}{\alpha_F^U}\right). \tag{S.130}$$

It is very nontrivial to sample from this probability distribution without introducing any intermediate variables, so we choose to apply the Metropolis-Hastings algorithm. First, Gamma distribution is selected as the proposal distribution, namely,

$$\left(\lambda_F^m\right)^{prop}\sim\textbf{Gamma}\left(\gamma_F^U,\frac{\left(\lambda_F^m\right)^{old}}{\gamma_F^U}\right). \tag{S.131}$$

The acceptance ratio of the proposed FRET rate then is

$$r_F^U = \frac{\mathbf{Gamma}\left((\lambda_F^m)^{old}; \gamma_F^U, \frac{(\lambda_F^m)^{prop}}{\gamma_F^U}\right) P\left((\lambda_F^m)^{prop} | d_{1:K}, \delta_{1:K}, c_{1:K}, \mathbf{x}_{1:K}, f_D, f_A, \lambda_E\right)}{\mathbf{Gamma}\left((\lambda_F^m)^{prop}; \gamma_F^U, \frac{(\lambda_F^m)^{old}}{\gamma_F^U}\right) P\left((\lambda_F^m)^{old} \middle| d_{1:K}, \delta_{1:K}, c_{1:K}, \mathbf{x}_{1:K}, f_D, f_A, \lambda_E\right)} \tag{S.132}$$

$$= \frac{\mathbf{Gamma}\left((\lambda_F^m)^{old}; \gamma_F^U, \frac{(\lambda_F^m)^{prop}}{\gamma_F^U}\right) \left[\prod_{\kappa_1=1}^{\mathcal{K}_1} P\left(\delta_{\kappa_1}, e_{\kappa_1} | (\lambda_F^m)^{prop}\right)\right]}{\mathbf{Gamma}\left((\lambda_F^m)^{prop}; \gamma_F^U, \frac{(\lambda_F^m)^{old}}{\gamma_F^U}\right) \left[\prod_{\kappa_1=1}^{\mathcal{K}_1} P\left(\delta_{\kappa_1}, e_{\kappa_1} \middle| (\lambda_F^m)^{old}\right)\right]} \times \frac{\mathbf{Gamma}\left((\lambda_F^m)^{prop}; \alpha_F^U, \frac{\beta_F^U}{\alpha_F^U}\right)}{\mathbf{Gamma}\left((\lambda_F^m)^{old}; \alpha_F^U, \frac{\beta_F^U}{\alpha_F^U}\right)}. \tag{S.133}$$

Note that

$$\frac{\mathbf{Gamma}\left((\lambda_F^m)^{old}; \gamma_F^U, \frac{(\lambda_F^m)^{prop}}{\gamma_F^U}\right) \mathbf{Gamma}\left((\lambda_F^m)^{prop}; \alpha_F^U, \frac{\beta_F^U}{\alpha_F^U}\right)}{\mathbf{Gamma}\left((\lambda_F^m)^{prop}; \gamma_F^U, \frac{(\lambda_F^m)^{old}}{\gamma_F^U}\right) \mathbf{Gamma}\left((\lambda_F^m)^{old}; \alpha_F^U, \frac{\beta_F^U}{\alpha_F^U}\right)}$$

$$= \frac{\left[(\lambda_F^m)^{prop}\right]^{\alpha_F^U - 2\gamma_F^U} \exp\left[-(\lambda_F^m)^{prop} \alpha_F^U/\beta_F^U + (\lambda_F^m)^{old} \alpha_F^U/\beta_F^U\right]}{\left[(\lambda_F^m)^{old}\right]^{\alpha_F^U - 2\gamma_F^U} \exp\left[-(\lambda_F^m)^{prop} \gamma_F^U/(\lambda_F^m)^{old} + (\lambda_F^m)^{old} \gamma_F^U/(\lambda_F^m)^{prop}\right]}, \tag{S.134}$$

the acceptance ratio gets simplified to

$$r_F^U = \frac{\prod_{\kappa_1=1}^{\mathcal{K}_1} P\left(\delta_{\kappa_1}, e_{\kappa_1} | (\lambda_F^m)^{prop}\right)}{\prod_{\kappa_1=1}^{\mathcal{K}_1} P\left(\delta_{\kappa_1}, e_{\kappa_1} \middle| (\lambda_F^m)^{old}\right)} \left[\frac{(\lambda_F^m)^{prop}}{(\lambda_F^m)^{old}}\right]^{\alpha_F^U - 2\gamma_F^U} \times \frac{\exp\left[-(\lambda_F^m)^{prop} \alpha_F^U/\beta_F^U + (\lambda_F^m)^{old} \alpha_F^U/\beta_F^U\right]}{\exp\left[-(\lambda_F^m)^{prop} \gamma_F^U/(\lambda_F^m)^{old} + (\lambda_F^m)^{old} \gamma_F^U/(\lambda_F^m)^{prop}\right]}. \tag{S.135}$$

#### C.8 Learning donor excitation rate

The corresponding conditional posterior is

$$P\left(\lambda_E \middle| d_{1:K}, c_{1:K}, \mathbf{x}_{1:K}, f_D, f_A, \lambda_F^{1:M}\right) \propto \left[\prod_{k=1}^{K} P\left(d_k \middle| c_k, \mathbf{x}_k, f_D, f_A, \lambda_E, \lambda_F^{1:M}\right)\right] P(\lambda_E) \tag{S.136}$$

$$\propto \left(\prod_{k=1}^{K} P_{d_k=\varnothing} p_{d_k=D}^{\delta_{1,d_k}} p_{d_k=A}^{\delta_{2,d_k}}\right) \mathbf{Gamma}\left(\lambda_E; \alpha_E, \frac{\beta_E}{\alpha_E}\right). \tag{S.137}$$

Again, we cannot sample from this probability distribution directly. Instead, we implement HMC once again.

From the negative log posterior, we can define the "potential energy" as

$$V(\lambda_E) = -\sum_{k=1}^{K} \left(\ln P_{d_k=\varnothing} + \delta_{1,d_k} \ln P_{d_k=D} + \delta_{2,d_k} \ln P_{d_k=A}\right) - (\alpha_E - 1) \ln \lambda_E + \frac{\alpha_E}{\beta_E}\lambda_E. \quad \text{(S.138)}$$

Then the full Hamiltonian is

$$\mathcal{H}\left(\lambda_E, p\right) = \frac{p^2}{2m} + V\left(\lambda_E\right). \quad \text{(S.139)}$$

We then evolve this system with the following steps:

1. advance a half-step using $V\left(\lambda_E\right)$,

$$p^{int} = p^i - \frac{h}{2}\frac{\partial V}{\partial \lambda_E}\bigg|_{(\lambda_E)^i}; \quad \text{(S.140)}$$

2. advance a whole-step using $T(p)$,

$$(\lambda_E)^f = (\lambda_E)^i + h\frac{p^{int}}{m}; \quad \text{(S.141)}$$

3. advance a half-step using $V\left(\lambda_E\right)$,

$$p^f = p^{int} - \frac{h}{2}\frac{\partial V}{\partial \lambda_E}\bigg|_{(\lambda_E)^f}. \quad \text{(S.142)}$$

All the needed derivatives are specified below

$$\frac{\partial V}{\partial \lambda_E} = -\frac{\partial}{\partial \lambda_E}\sum_{k=1}^{K} \left(\ln P_{d_k=\varnothing} + \delta_{1,d_k} \ln P_{d_k=D} + \delta_{2,d_k} \ln P_{d_k=A}\right) - \frac{\alpha_E - 1}{\lambda_E} + \frac{\alpha_E}{\beta_E} \quad \text{(S.143)}$$

$$= -\sum_{k=1}^{K}\left(\frac{1}{P_{d_k=\varnothing}}\frac{\partial P_{d_k=\varnothing}}{\partial \lambda_E} + \frac{\delta_{1,d_k}}{p_{d_k=D}}\frac{\partial p_{d_k=D}}{\partial \lambda_E} + \frac{\delta_{2,d_k}}{p_{d_k=A}}\frac{\partial P_{d_k=A}}{\partial \lambda_E}\right) - \frac{\alpha_E - 1}{\lambda_E} + \frac{\alpha_E}{\beta_E}. \quad \text{(S.144)}$$

$$\frac{\partial P_{d_k=\varnothing}}{\partial \lambda_E} = \frac{\partial}{\partial \lambda_E}\left(e^{\mu^D_{E,k}\tau} + e^{\mu^A_{E,k}\tau} + e^{\lambda^D_B T} + e^{\lambda^A_B T} - 3\right)^{-1} \quad \text{(S.145)}$$

$$= -\tau \mathrm{PSF}_k P^2_{d_k=\varnothing}\left(f_D e^{\mu^D_{E,k}\tau} + f_A \xi e^{\mu^A_{E,k}\tau}\right), \quad \text{(S.146)}$$

$$\frac{\partial p_{d_k=D}}{\partial \lambda_E} = \tau \mathrm{PSF}_k\left(f_D e^{\mu^D_{E,k}\tau}\frac{\mu^D_R P_{DD} + \mu^{c_k}_F P_{AD}}{\mu^D_R + \mu^{c_k}_F} + f_A \xi e^{\mu^A_{E,k}\tau} P_{AD}\right), \quad \text{(S.147)}$$

$$\frac{\partial p_{d_k=A}}{\partial \lambda_E} = \tau \mathrm{PSF}_k\left(f_D e^{\mu^D_{E,k}\tau}\frac{\mu^D_R P_{DA} + \mu^{c_k}_F P_{AA}}{\mu^D_R + \mu^{c_k}_F} + f_A \xi e^{\mu^A_{E,k}\tau} P_{AA}\right). \quad \text{(S.148)}$$

### 4 Supplementary tables

Table S.1: List of abbreviations.

| Phrase | Abbreviation |
|---|---|
| Fluorescence correlation spectroscopy | FCS |
| Region of interest | ROI |
| Hamiltonian Monte Carlo | HMC |
| Point spread function | PSF |
| Markov chain Monte Carlo | MCMC |
| Maximum a posteriori | MAP |
| Instrument response function | IRF |

Table S.2: List of calibrated parameters used in data analysis.

| Quantity | Value |
|---|---|
| PSF width in $x$ | 0.3 µm |
| PSF width in $y$ | 0.3 µm |
| PSF width in $z$ | 1.5 µm |
| Interpulse interval | 51.37 ns |
| Pulse width | 10 ps |
| Acceptor direct excitation ratio ($\xi$) | 0.05 |
| IRF mean | 1.26 ns |
| IRF standard deviation | 0.388 ns |
| Donor channel background photon rate | $2 \times 10^3\,s^{-1}$ |
| Acceptor channel background photon rate | $2 \times 10^3\,s^{-1}$ |
| Donor-to-acceptor bleed-through probability | 0.06 |
| Acceptor-to-donor bleed-through probability | 0.01 |

Table S.3: Probability distributions used and their densities. Here, the corresponding random variables are denoted by $x$.

| Distribution | Notation | Probability density function | Mean value | Variance Covariance |
|---|---|---|---|---|
| Normal | $\mathbf{Normal}(\mu, \sigma^2)$ | $\frac{1}{\sqrt{2\pi\sigma^2}} e^{-\frac{(x-\mu)^2}{2\sigma^2}}$ | $\mu$ | $\sigma^2$ |
| Exponential | $\mathbf{Exponential}(\tau)$ | $\frac{1}{\tau} e^{-x/\tau}$ | $\tau$ | $\tau^2$ |
| Gamma | $\mathbf{Gamma}(\alpha, \beta)$ | $\frac{1}{\Gamma(\alpha)\beta^\alpha} x^{\alpha-1} e^{-\frac{x}{\beta}}$ | $\alpha\beta$ | $\alpha\beta^2$ |
| Inverse Gamma | $\mathbf{InvGamma}(\alpha, \beta)$ | $\frac{\beta^\alpha}{\Gamma(\alpha)} x^{-\alpha-1} e^{-\frac{\beta}{x}}$ | $\frac{\beta}{\alpha-1}$ | $\frac{\beta^2}{(\alpha-1)^2(\alpha-2)}$ |
| Beta | $\mathbf{Beta}(\alpha, \beta)$ | $\frac{\Gamma(\alpha+\beta)}{\Gamma(\alpha)\Gamma(\beta)} x^{\alpha-1}(1-x)^{\beta-1}$ | $\frac{\alpha}{\alpha+\beta}$ | $\frac{\alpha\beta}{(\alpha+\beta)^2(\alpha+\beta+1)}$ |
| Dirichlet | $\mathbf{Dirichlet}(\alpha, \beta)$ | $\frac{\Gamma(\alpha+\beta)}{\Gamma(\alpha)\Gamma(\beta)} x^{\alpha-1}(1-x)^{\beta-1}$ | $\frac{\alpha}{\alpha+\beta}$ | $\frac{\alpha\beta}{(\alpha+\beta)^2(\alpha+\beta+1)}$ |
| Bernoulli | $\mathbf{Bernoulli}(p)$ | $(1-p)^{\delta_0(x)} p^{\delta_1(x)}$ | $q$ | $q(1-q)$ |
| Categorical | $\mathbf{Categorical}(p_{1:M})$ | $\prod_{m=1}^{M} p_m^{\delta_m(x)}$ | | |
